# Differential chromatin looping regulated by two GA-binding transcription factors creates an X-specific chromatin environment for dosage compensation

**DOI:** 10.64898/2026.03.12.711449

**Authors:** Joseph Aguilera, Kaitlyn Cortez, Luna C. Segarra Alonzo, Melissa Aldana, Angelica Aragon, Claire Gray, Morgan Woodman-Sousa, Kathryn J. Grive, Mukulika Ray, Erica Larschan

**Affiliations:** Department of Molecular Biology, Cell Biology, and Biochemistry, Brown University, Providence, RI; Department of Biology, University of Puerto Rico, Río Piedras, San Juan, PR; Department of Pharmacology and Cancer Biology, Duke University, Durham, NC; Department of Human Genetics, University of Chicago, Chicago, IL; Department of Biology, University of Massachusetts, Boston, Boston, MA; Women and Infants Hospital of Rhode Island, Department of Obstetrics and Gynecology, Program in Women’s Oncology, Providence, RI

## Abstract

The mechanisms by which differential occupancy of transcription factors (TFs) at similar binding sites leads to context-specific targeting of large transcription complexes remain poorly understood. X chromosome upregulation (XCU), the most highly conserved step in dosage compensation and best studied in *Drosophila*, serves as a model for understanding how differential occupancy of similar TFs functions context-specifically. Sequence variation within GA-repeat motifs that accumulated on the X chromosome over evolutionary time promotes the binding of a specific GA-binding TF (CLAMP) that recruits the dosage compensation complex (DCC) while outcompeting another similar TF (GAF). However, the mechanism by which CLAMP-GAF competition drives specific targeting of the DCC to the X chromosome remains unknown. Because DCC binding sites cluster in 3D space, we combined Micro-C and Hi-ChIP to determine that CLAMP and GAF directly mediate largely mutually exclusive 3D genomic contacts. Specifically, we show that CLAMP but not GAF drives local short-range interactions that directly link high affinity DCC binding sites with active, dosage-compensated housekeeping genes. In contrast, GAF mediates interactions between transcriptionally silent insulator regions on the X chromosome spanning a wider range of genomic distances. Together, these findings demonstrate that CLAMP outcompetes GAF at active regions on the X chromosome, but not autosomes, to create an X-chromosome specific chromatin environment for dosage compensation. Overall, we provide new insight into how differential TF binding at similar binding sites drives context-specific targeting of transcription complexes.

## Introduction

Within the nucleus of eukaryotic cells, chromatin is tightly compacted through multiple orders of organization (Bonev and Cavalli 2016) and shares finite nuclear space with many classes of transcriptional regulators, including DNA-binding transcription factors (TFs) and large transcription complexes which are often targeted to chromatin by TFs (Zaret 2020). TFs recruit large transcription complexes to facilitate spatially precise and temporally dynamic transcriptional repression and activation of specific groups of genes (Zaret 2020; Drouin 2014; Wu 1995). Furthermore, TFs such as CTCF are important for defining boundaries between three dimensional domains by regulating the loop extrusion process among other functions (Barth et al. 2025) (Merkenschlager and Nora 2016). In mammals, additional factors that bind to similar binding sites function redundantly with CTCF but their relationship with CTCF remains poorly understood (Skok et al. 2022).

In *Drosophila*, TFs and transcription complexes play an important role facilitating chromatin loops that bring together specific groups of genes for organization and transcription through the function of insulators and tethering elements (Gutierrez-Perez et al. 2019; Boettiger et al. 2016; Melnikova et al. 2004; Wolle et al. 2015; Kaye et al. 2017; Li et al. 2023; Varisco et al. 2025; Levo et al. 2022). Furthermore, recent work indicates that loop extrusion mechanisms function together with tethering elements containing GA-rich sequences to promote loop formation and transcriptional bursting (Choppakatla et al. 2026). While loop extrusion increases the rate of the search for the correct 3D contact, tethering elements are essential to stabilize these contacts and promote transcriptional bursting through heterotypic interactions between different TFs (Varisco et al. 2025; Choppakatla et al. 2026). However, the mechanisms by which specific TFs bring together the correct target loci in a context-specific manner to precisely coregulate genes remain unknown across species.

To understand how specific TFs mediate chromatin loops context-specifically to drive specific transcriptional programs, it is important to use a model in which: 1) specific chromatin loops mediate a particular transcriptional program; 2) a specific TF has been identified that regulates the transcriptional program excluding the function of another known TF. Many TFs that are members of similar functional classes occupy similar binding sites (Zaret 2020). Furthermore, variation within these sites, including SNPs associated with disease, can drive differential occupancy of TFs to similar binding sites (Jeong and Bulyk 2025; Khetan et al. 2025; Jaeger et al. 2010; Kuzu et al. 2016). Yet, in most contexts little is known about the differential functions of these specific TFs in mediating different transcriptional programs or their direct function in regulating 3D loop formation.

Therefore, we used the established *Drosophila* X-chromosome upregulation (XCU) dosage compensation system in which the presence of a specific GA-binding TF has a direct role in mediating the recruitment of the dosage compensation complex (DCC) (Soruco et al. 2013; Larschan et al. 2012; Eggers et al. 2023; Eggers and Becker 2021) compared with another similar factor (Kaye et al. 2018). XCU, which equalizes the levels of X-chromosome and autosomal gene expression within heterogametic species, is the most highly conserved step in dosage compensation across species that maintains the stoichiometry of the many protein complexes with components encoded on both the X chromosome and autosomes (Samata and Akhtar 2018). Recent work has identified convergent evolution toward the enrichment of the H4K16ac chromatin mark which increases transcript levels two-fold as the most highly conserved mechanism for dosage compensation which occurs across heterogametic species (Zimmer et al. 2025; Gu et al. 2019). Because XCU was first identified in *Drosophila,* which has excellent tools for dissecting transcriptional mechanisms, it has been best studied in this system (Samata and Akhtar 2018).

Variation within GA-rich binding sites that accumulated over evolutionary time on the X chromosome promotes the occupancy of the pioneer TF CLAMP (Chromatin Linked Adaptor for MSL Proteins) instead of the well-studied pioneer TF GAGA-Factor (GAF) (Kaye et al. 2018; Kuzu et al. 2016). CLAMP mediates DCC binding to the X chromosome to facilitate the two-fold upregulation of the single male X chromosome (Soruco et al. 2013; Eggers et al. 2023). Apart from dosage compensation, these two TFs have opposing transcriptional functions at most genomic loci: GAF often acts as a transcriptional activator (Adkins et al. 2006; Duarte et al. 2016), while CLAMP often functions as a repressor (Urban et al. 2017a). Therefore, understanding the competitive relationship between GAF and CLAMP will also reveal new insight into how the balance between activating and repressive transcriptional programs is established across the genome.

Despite decades of study, the mechanisms by which an X-chromosome-specific chromatin environment is established to recruit the DCC to the bodies of active genes remains poorly understood (Larschan et al. 2007; Alekseyenko et al. 2012). CLAMP binds to GA-rich sequence motifs on the X chromosome within the strongest DCC binding sites which are known as chromatin entry sites (CES) (Kelley et al. 1999) or high-affinity sites (HAS) (Straub et al. 2008). Subsequent to identifying these binding sites that are already occupied by CLAMP in early development (Rieder et al. 2019), the DCC spreads along the length of the X chromosome through the function of DCC components, including the *roX* non-coding RNAs (Rieder et al. 2019; Valsecchi et al. 2020; Park et al. 2002; Kelley et al. 1999).

Beyond its function in XCU, CLAMP is a pioneer TF that binds to GA-rich motifs to regulate chromatin accessibility and remodeling in other contexts such as zygotic genome activation (Duan et al. 2021) and histone gene transcription (Rieder et al. 2017). GAF is one of the first TFs to be identified across species and recognizes GA-rich motifs throughout the genome to mediate activation of transcription in many contexts including zygotic genome activation (Gaskill et al. 2021) and regulation of the heat shock genes (Duarte et al. 2016; Adkins et al. 2006). Variation in sequence composition at GA-motifs determines whether CLAMP or GAF will occupy a specific site, with CLAMP but not GAF tolerating variation with stretches of GA-repeats that occurs within the strongest DCC binding sites (HAS/CES) (Kaye et al. 2018; Adkins et al. 2006). While CLAMP but not GAF directly associates with the DCC both *in vivo* and *in vitro* (Alekseyenko et al. 2015; Eggers et al. 2023), GAF promotes DCC recruitment but to a much lesser extent (Kaye et al. 2018). Therefore, XCU serves as a model for understanding how small differences in DNA sequence determine the context-specific recruitment of large transcription complexes that induce coordinated regulation of specific groups of genes.

Prior low-resolution studies suggested that the strongest DCC binding sites cluster in 3D space and CLAMP contributes to this process (Jordan and Larschan 2021). GAF has also been shown to contribute to 3D chromatin looping in the nervous system (Li et al. 2023). Therefore, we hypothesized that the competition between CLAMP and GAF leads to context-specific formation of chromatin looping that differs on the X chromosome compared to autosomes. To define how factors with similar binding sites differentially regulate chromatin loop formation, we determined the individual direct functions of GAF and CLAMP on the X chromosome and autosomes by conducting both Micro-C and Hi-ChIP. We first identified CLAMP and GAF binding sites using Hi-ChIP, then identified differential chromatin looping within the nucleus after knocking down GAF and CLAMP individually using Micro-C. By integrating Hi-ChIP and Micro-C we define direct roles that CLAMP and GAF play in chromatin loop formation at up to 1kb resolution.

Specifically, we demonstrate that the differential occupancy of CLAMP versus GAF which is caused by minor sequence variation within GA-rich motifs on the X chromosome(Kaye et al. 2018) drives the formation of local short-range interactions that can directly link high affinity DCC binding sites with active housekeeping genes that are dosage compensated. In contrast, GA-repeat motifs that are contiguous promote GAF binding that can drive interactions between silent insulator regions on the X chromosome and span a broad range of genomic distances. On autosomes, both CLAMP and GAF directly regulate contacts between active chromatin regions including active promoters. Therefore, CLAMP outcompetes GAF at active regions on the X chromosome but not autosomes to mediate the short-range 3D contacts required for DCC recruitment specifically to active genes on the X-chromosome that need to be dosage compensated. CLAMP but not GAF can directly interact with the DCC to recruit it to its target genes where it upregulates transcription (Eggers et al. 2023; Eggers and Becker 2021), suggesting that these 3D contacts are important for DCC occupancy (Soruco et al. 2013) and transcriptional upregulation (Urban et al. 2017a) that are both mediated by CLAMP. Overall, we provide key insight into how differential binding of two essential TFs to similar binding sites drives context-specific targeting of a transcription complex.

## Results

### 1. CLAMP- and GAF-dependent chromatin loops are largely non-overlapping and have different size distributions

To compare how CLAMP and GAF regulate chromatin loop formation, we used *Drosophila* S2 cells as a model system for the following reasons. S2 cells are embryonically derived male *Drosophila* cells that have been used to study XCU for several decades (Straub et al. 2008; Alekseyenko et al. 2012, 2008; Valsecchi et al. 2020). The DCC binds to very similar loci in S2 cells compared with embryos and neuronally derived cell lines. This suggests that using S2 cells provides sufficient biological context while also circumventing asynchronous embryonic lethality that occurs after the loss of maternally derived CLAMP in embryos (Rieder et al. 2017; Alekseyenko et al. 2008, 2012).

To define the function of the CLAMP and GAF GA-binding transcription factors in shaping the 3D chromatin architecture of the *Drosophila* genome, we utilized a high-resolution chromatin mapping technique, Micro-C (Hsieh et al. 2015), with established RNAi-mediated knockdown approaches (Soruco et al. 2013; Larschan et al. 2012; Kaye et al. 2018) in *Drosophila* S2 cells (Fig. 1A). We validated *clamp* and *gaf* RNAi knockdowns using Western blotting (Fig. S1). Whole-chromosome differential contact maps revealed that depletion of either CLAMP or GAF disrupts genome-wide looping, substantially remodeling the 3D architecture of the *Drosophila* genome (Fig. S2).

**Figure 1.**
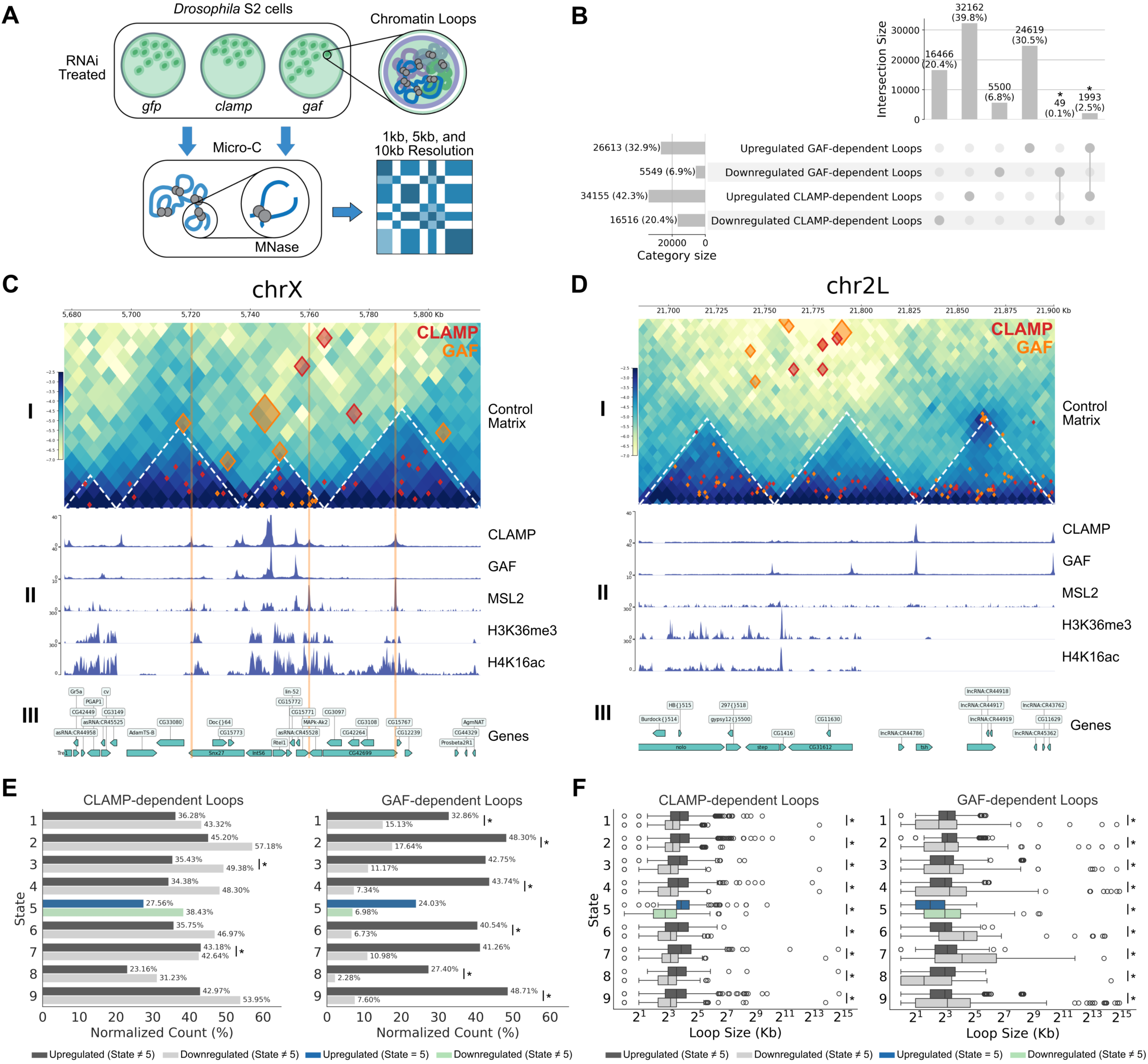
Loss of CLAMP versus GAF alters different chromatin loops. **(A)** Experimental workflow for Micro-C analysis. Schematic illustrating RNAi-mediated depletion of CLAMP or GAF in *Drosophila* S2 cells, with *gfp* RNAi serving as a control. Micro-C libraries were generated using MNase digestion and sequenced at 1kb, 5kb, and 10kb resolutions. **(B) Loops altered by loss of CLAMP or GAF overlap at a low frequency.** UpSet plot showing the intersection of significantly altered chromatin loops (Fisher’s Exact Test, *P* < 0.05) following *clamp* RNAi (*clampi)* or *gaf* RNAi (*gafi)* treatment. Vertical bars represent the number of loops that are either upregulated (gained) or downregulated (lost) across the indicated categories. **(C-D) Representative Micro-C contact maps and genomic tracks.** CoolBox plots displaying the chromatin landscape on **(C)** the X chromosome (chrX) and **(D)** an autosome (chr2L). **(I)** Control Micro-C contact matrices (*gfpi*) showing differential CLAMP-(red diamonds) and GAF-dependent (orange diamonds) loops. Size of diamond corresponds to the resolution of the differential chromatin interaction: small (1kb), medium (5kb), and large (10kb). **(II)** Normalized ChIP-seq tracks for CLAMP, GAF, MSL2, H3K36me3, and H4K16ac across a representative 200 kb genomic window. **(III)** *Drosophila* (*dm6*) genes are shown at the bottom; boxes represent exons and arrows indicate transcriptional orientation. **(E) Chromatin state enrichment of differential loops.** Normalized counts (%) of CLAMP- and GAF-dependent loops categorized by the 9-state chromatin model. Asterisks (*) indicate significant difference in enrichment (Fisher’s Exact Test, *P* < 0.05). **(F) Genomic length span of state-specific loops.** Boxplots showing the loop size (kb) distribution for CLAMP- and GAF-dependent loops across chromatin states 1-9. Asterisks (*) indicate significant difference between loop sizes (Mann-Whitney Test, *P* < 0.05). For panels **E** and **F**, upregulated loops in State 5 (dosage compensation) are highlighted in blue, while downregulated State 5 (dosage compensation) loops are highlighted in green.

The depletion of the GA-binding transcription factors CLAMP and GAF profoundly disrupted higher-order chromatin organization. We first identified 3,852 TADs in control conditions (Fig. S3A). Depletion of CLAMP altered 1,218 (31.62%) of these TADs, while GAF depletion had an even more dramatic effect, disrupting 3,415 (88.66%) of all TADs (Fig. S3A, C). Notably, 1,044 of these domains, representing 27.1% of all control TADs, were disrupted in both knockdowns, a highly significant overlap (*P* < 0.001, Fisher’s Exact Test) (Fig. S3B). The widespread loss of TAD integrity after CLAMP and GAF depletion underscores the critical role of these TFs in maintaining chromatin architecture.

To maximize the high-resolution capabilities of Micro-C, we performed differential chromatin loop analyses at three resolutions (1kb, 5kb, and 10kb) for each knockdown condition (*clamp* or *gaf* RNAi) to detect both short-and long-range differential loops (Fig. 1A). We integrated the multi-resolution loops for each condition which we then used throughout our analysis (Table S1).

Differential loop quantification with the HiCcompare R-package (Stansfield et al. 2018) revealed four distinct sets of loops (Fig. 1B): 1) chromatin loops that are **upregulated** after *clamp* RNAi relative to control *gfp* RNAi (*upregulated CLAMP-dependent loops*, N=34,155); 2) chromatin loops that are **downregulated** after *clamp* RNAi relative to control *gfp* RNAi (*downregulated CLAMP-dependent loops*, N=16,516); 3) chromatin loops that are **upregulated** after *gaf* RNAi relative to control *gfp* RNAi *(upregulated GAF-dependent loops*, N=26,613); 4) chromatin loops that are **downregulated** after control *gaf* RNAi relative to control *gfp* RNAi (*downregulated GAF-dependent loops*, N=5,549). Across the four sets of differential chromatin loops, we only observed very minimal direct overlap between the following groups (Fig. 1B): upregulated CLAMP- and GAF-dependent loops (49 loops; 0.1%; *P* < 0.01, Fisher’s Exact Test) and downregulated CLAMP- and GAF-dependent loops (1,993 loops, 2.5%, *P* < 0.01, Fisher’s Exact Test). The minimal direct overlap observed between differential CLAMP- and GAF-dependent loops suggests that CLAMP and GAF have largely mutually exclusive looping roles in the genome even though they sometimes regulate loops within the same TADs.

Interestingly, CLAMP depletion resulted in far fewer differential TADs than GAF despite yielding substantially more differential loops (Fig.1B, Fig. S3). Furthermore, 85.71% of TADs disrupted by CLAMP knockdown were also disrupted by GAF knockdown, whereas only 30.57% of GAF-disrupted TADs overlapped with those affected by CLAMP depletion (Fig. S3). These data suggest that GAF may have a more prominent role in maintaining TAD integrity than CLAMP. The observation that CLAMP depletion produces more differential loops, yet fewer disrupted TADs, implies that CLAMP may primarily regulate precise chromatin looping within TADs, whereas GAF is more critical for establishing or maintaining the broader domain boundaries that define TAD structure.

Subsequently, we performed a comparative visualization of a 220.5kb X-chromosome region (chrX:5,604,766-5,825,266) and an 185kb autosomal region (chr2L:21,660,000-21,845,000). For each region, we examined differential chromatin loops relative to local TADs, CLAMP and GAF occupancy, MSL2 (the DCC core protein subunit), histone modifications, and the *Drosophila* S2 9-state chromatin model (Fig. 1C, D).

A key looping characteristic emerged through visualization in both genomic contexts, revealing a resolution-dependent distinction between intra-TAD and inter-TAD loop organization. On the X chromosome, we first visualized all differential loops over the control (*gfp* RNAi) Micro-C contact matrix and its corresponding TADs, with the region’s High Affinity Sites (HAS) highlighted in orange to contextualize the 3D chromatin changes and factor occupancy (Fig. 1C: I). With this visualization established, we observed that high-resolution (1kb) differential loops dependent on CLAMP and GAF were primarily contained within TADs, while lower-resolution (5kb and 10kb) loops often connected regions outside of these domains. Notably, this same resolution-dependent characteristic was also present on the autosomal region, where 1kb loops were intra-TAD and lower-resolution loops extended to inter-TAD interactions (Fig. 1D: I). This consistent resolution-dependent partitioning of intra- and inter-TAD loops underscores that multi-resolution analysis is essential to capture the complete and diverse set of architectural changes mediated by these factors, regardless of the chromosome.

When comparing CLAMP and GAF occupancy at the X-chromosome region, our findings are consistent with prior findings that CLAMP outcompetes GAF at HAS which has been demonstrated *in vivo* (Kaye et al. 2018) and *in vitro* (Eggers and Becker 2021; Eggers et al. 2023). CLAMP is enriched while GAF is absent in the highlighted HAS (Fig. 1C: II). Importantly, MSL2 is highly enriched at the HAS, consistent with findings from previous studies which outline the synergistic relationship between CLAMP and MSL2 at the HAS (Eggers et al. 2023; Soruco et al. 2013). To mark DCC activity, we observed the presence of H4K16ac (Gu et al. 1998) and visualized the H3K36me3 chromatin mark to show the presence of active gene bodies (Larschan et al. 2007) in this region (Fig. 1C: II, III). Importantly, these histone modifications are enriched flanking the HAS, consistent with local spreading of the DCC from its initialization sites (Kelley et al. 1999) visualized in two dimensions. Conversely, the autosomal region lacks both HAS and MSL2 enrichment as expected (Fig. 1D: II, III). While CLAMP and GAF occupy the GA-repeats present throughout the genome, the lack of MSL2 and H4K16ac occupancy on the autosome region underscores the X-specificity of DCC recruitment.

After visualizing differential CLAMP- and GAF-dependent loops over specific candidate regions, we measured how these loops are distributed across different chromatin states globally using the *Drosophila* S2 9-state HMM chromatin model, which bins the genome into one of nine chromatin states by integrating dozens of modENCODE chromatin profiling data sets (Kharchenko et al. 2010). Importantly, we exclusively used differential loops detected at a 1kb resolution for the chromatin state enrichment analysis, as lower resolutions contained too much chromatin state diversity within loop anchors. For each set of differential CLAMP- and GAF-dependent loops, we determined whether there was a difference in chromatin state enrichment between upregulated and downregulated loops. Interestingly, differential CLAMP-dependent loops yielded only two significant enrichment differences, in the open chromatin (state 4) and heterochromatin (state 7), while differential GAF-dependent loops yielded significant enrichment differences across a broader range of chromatin environments, spanning active transcription (states 1 and 2), open chromatin (state 4), Polycomb-mediated repression (state 6), heterochromatin-like regions (state 8) and transcriptionally silent euchromatin (state 9) (Fig. 1E). The stark difference in chromatin state enrichment between CLAMP- and GAF-dependent loops, with only the open chromatin state (state 4) shared between them, suggests that these similar GA-binding TFs may regulate functionally distinct chromatin environments across the genome.

After measuring chromatin state enrichment, we asked whether the sizes of differential CLAMP- and GAF-dependent loops differ across chromatin states (Fig. 1F). Notably, within every chromatin state category, the size distributions of upregulated and downregulated loops were significantly different for both CLAMP and GAF knockdowns (Fig. 1F; *P* < 0.001, Mann-Whitney Test). However, in addition to this shared significance, a distinct TF-specific shift in the interquartile ranges (IQRs) emerged. When examining differential CLAMP-dependent loops, the IQRs of upregulated loops expanded and shifted upward relative to downregulated loops, indicating a loss of shorter-range interactions upon CLAMP depletion and a gain of longer-range interactions. In contrast, the opposite pattern was observed for differential GAF-dependent loops: the IQRs of the upregulated loops contracted and shifted downward relative to downregulated loops, indicating a loss of longer-range interactions upon GAF depletion and a gain of shorter-range interactions. Together, these opposing size shifts suggest that CLAMP and GAF have different effects on chromatin looping: CLAMP loss preferentially disrupts local interactions, whereas GAF loss disrupts longer range interactions.

Importantly, the global looping characteristics described above for CLAMP- and GAF-dependent loops also occurred at dosage compensated genes (state 5) (Fig. 1E, F). Given that *Drosophila* dosage compensation is regulated by well-characterized DNA regulatory elements (Alekseyenko et al. 2008, 2012), TF competition (Kaye et al. 2018), and a large chromatin modifying complex (Samata and Akhtar 2018), we can leverage this conserved biological process (Zimmer et al. 2025; Deng et al. 2013) to understand how GAF and CLAMP function differentially to regulate transcription context-specifically.

### 2. CLAMP and GAF are required for different sized chromatin loops near GA-repeat rich high-affinity sites (HAS)

Due to the established competition between CLAMP and GAF high affinity sites (HAS) that regulates DCC recruitment (Kaye et al. 2018), we next examined how CLAMP and GAF regulate chromatin looping specifically at HAS. We quantified the proportion of dosage-compensated genes contained within the differential CLAMP-and GAF-dependent loops, and which were also associated with the HAS. We determined that CLAMP-dependent loops encompass over 90% of dosage-compensated (DC) genes anchored at HAS on the X chromosome, whereas GAF-dependent loops only encompassed 50% or fewer DC genes associated with HAS (Fig. 2A). Furthermore, CLAMP-dependent loops were found to overlap the HAS more frequently than GAF-dependent loops (Fig. 2A). These data are consistent with the established stronger requirement for CLAMP in regulating DCC recruitment to HAS compared with GAF which also has a significant although smaller role which had not yet been defined (Kaye et al. 2018).

**Figure 2.**
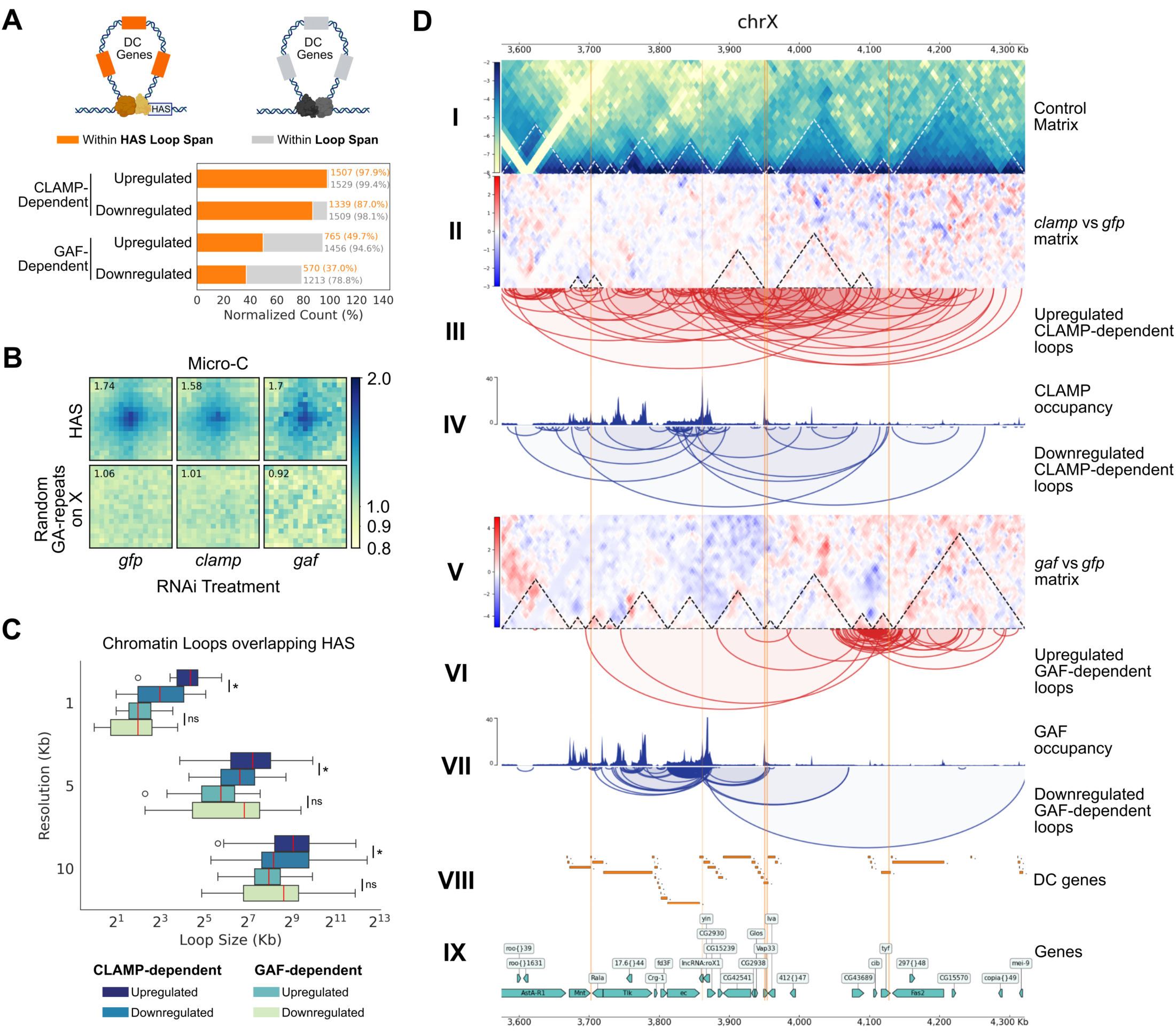
Loss of CLAMP and GAF alters chromatin loop architecture at High Affinity Sites (HAS) on the X chromosome. **(A)** Association of CLAMP- and GAF-dependent loops with dosage-compensated (DC) genes. Top: Schematic of: **Left:** DC genes (orange) localized within the span of loops anchored at High Affinity Sites (HAS). **Right:** DC genes not within HAS loop span. **Bottom:** Normalized counts (%) of upregulated and downregulated CLAMP- and GAF-dependent loops containing DC genes within their span. Numbers indicate the total DC gene count and the percentage of DC genes that are encompassed by loops either associated with HAS or not. **(B) Aggregate Peak Analysis (APA) of HAS contacts.** Heatmaps showing the average Micro-C contact enrichment centered on High Affinity Sites (top row) compared to random GA-repeat sequences on the X chromosome (bottom row). Enrichment scores (upper left) indicate a reduction in contact strength following *clamp* RNAi (enrichment = 1.58) or *gaf* RNAi (enrichment = 1.7) treatment compared to *gfp* RNAi control (enrichment = 1.74). Pixel resolution is 2.5 kb with ± 20kb flanking central pixel. **(C) Distribution of loop sizes at HAS.** Box plots categorized by Micro-C resolution (1kb, 5kb, 10kb) showing the genomic span (Kb) of upregulated and downregulated CLAMP- and GAF-dependent loops overlapping HAS. **(D) Representative chromatin loops regulated by CLAMP or GAF on the X chromosome at HAS (CoolBox visualization) (I)** Control Micro-C contact matrix (*gfpi*). **(II & V)** Differential contact matrices showing log_2_ fold-changes for *clampi* vs *gfpi* and *gafi* vs *gfpi*. Red indicates increased contacts and blue indicates decreased contacts. Differential TADs are shown with black dashed lines. (**III, IV, VI, VII)** Loop tracks representing upregulated (red) and downregulated (blue) loops for CLAMP and GAF, respectively, aligned with their corresponding protein occupancy Hi-ChIP tracks. **(VIII)** Genomic tracks for DC genes and (IX) *dm6* gene annotations.

Therefore, we dissected the differences between GAF and CLAMP regulated loops at HAS because the loops they regulate encompass different percentages of DC genes. Micro-C APA plots revealed robust looping at the HAS (log enrichment = 1.74) that diminished substantially upon CLAMP knockdown (log enrichment = 1.58), while GAF depletion caused a more diffuse loss of signal distal at the HAS (log enrichment = 1.70) (Fig. 2B).

Additionally, we observed a distinct relationship between loop sizes: After CLAMP knockdown, downregulated loops over HAS were considerably smaller than upregulated loops (Fig. 2C; *P* < 0.05, Mann-Whitney Test). The 1kb resolution provided by Micro-C reveals CLAMP anchors precisely at the center of HAS at the nucleosomal-scale level (Fig. 2C; *P* < 0.001; Mann-Whitney Test) that may have otherwise been blurred at 10kb or masked by multi-resolution aggregation (Fig. 2C; *P* < 0.05; Mann-Whitney Test). Conversely, GAF knockdown loop distribution over HAS showed that downregulated loops maintained a larger median loop size than upregulated loops (Fig. 2C). This variation in loop size was best observed at 5kb and 10kb resolutions, where larger loops can be captured (Fig. 2C). These findings suggest that CLAMP and GAF may maintain distinct, size-specific regulatory roles at the HAS, an observation strongly correlated with the specific loop size ranges identified by Micro-C.

Examination of a 747 kb candidate region on the X chromosome (chrX:3,575,000-4,322,000) further illustrated the divergent effects of CLAMP and GAF loss on chromatin looping near HAS (Fig. 2D). The increase in contacts surrounding and within differential TADs upon CLAMP knockdown demonstrates that CLAMP is necessary for maintaining short, high-frequency loops at HAS (Fig. 2D: I, II, IV). Conversely, the loss of larger chromatin loops around differential TADs upon GAF knockdown demonstrates that GAF regulates the broader neighborhood surrounding HAS rather than only the specific HAS locus (Fig. 2D: I, V, VII). Notably, CLAMP depletion resulted in approximately twice as many upregulated than downregulated chromatin loops, a pattern visible within the candidate region (Fig. 2D: III). This asymmetry was even more pronounced upon GAF depletion, where upregulated loops outnumbered downregulated loops by approximately five-fold (Fig. 2D: VI). Together, our data support a model in which CLAMP and GAF function at HAS to cluster DC genes into 3D domains of different sizes (Fig. 2D: VIII, IX). However, it is critical to determine which loops are directly versus indirectly regulated by CLAMP and GAF to define their mechanism of action.

### 3. Single nucleotide variation within GA-rich motifs correlates with TF-specific chromatin looping at HAS

We observed that CLAMP and GAF influence loops of different sizes at HAS and throughout the genome (Fig. 1F, Fig. 2C). However, with Micro-C alone, we were not able to determine which differential loops were a result of a direct absence of either TF or an indirect effect mediated by other regulators. Therefore, we incorporated the MNase Hi-ChIP technique (Mumbach et al. 2016) targeting CLAMP and GAF to annotate the Micro-C differential loops to distinguish direct from indirect effects of TF-specific knockdowns (Fig. 3A). Notably, direct and indirect effects were only determined for the downregulated CLAMP- and GAF-dependent loops as these are the interactions that are present under control conditions which are the same conditions we used for Hi-ChIP.

**Figure 3.**
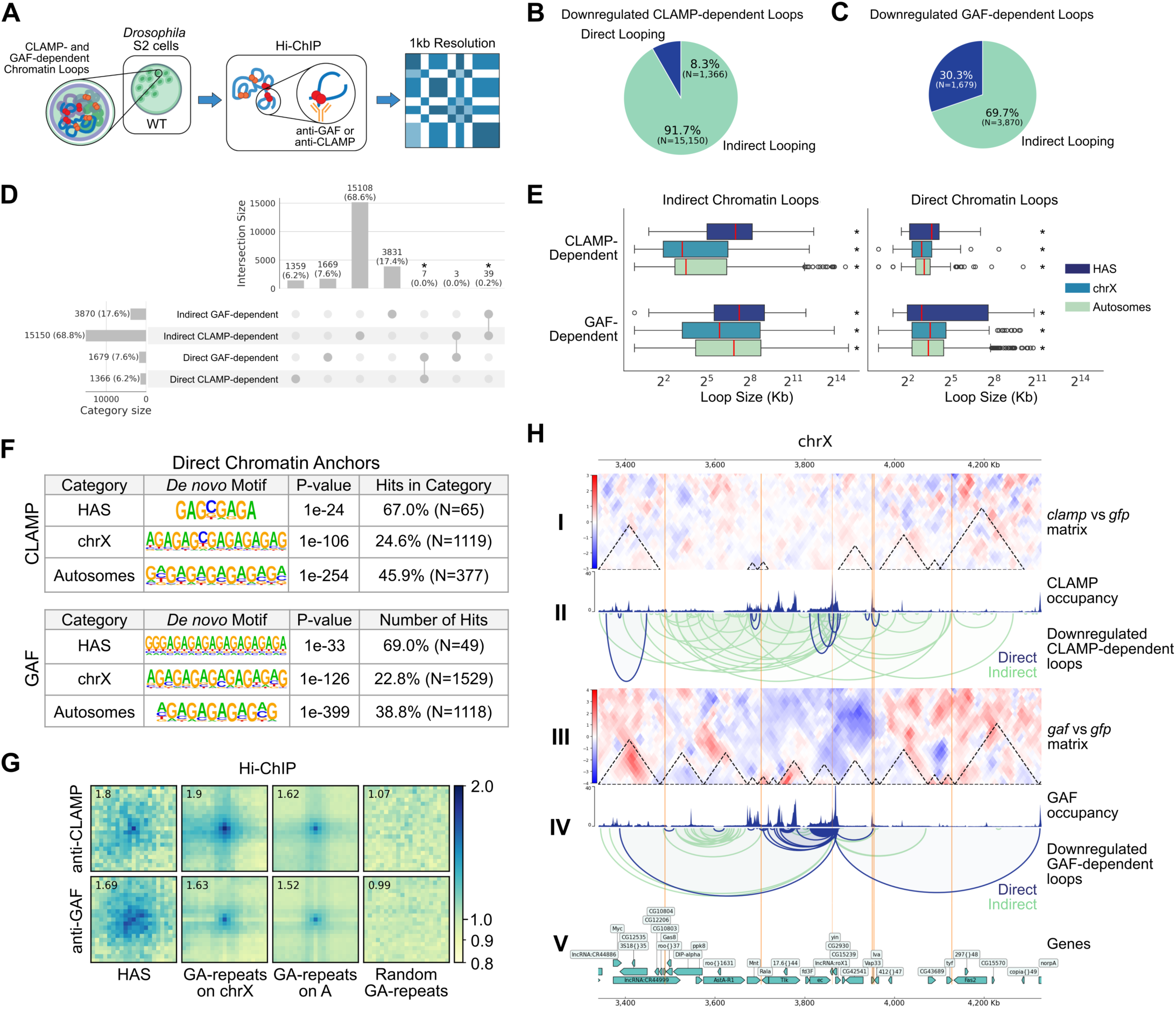
Direct looping by CLAMP and GAF suggests size-specific loop regulation at the HAS. **(A)** Hi-ChIP experimental workflow. Schematic illustrating the Hi-ChIP protocol in *Drosophila* S2 cells using anti-CLAMP and anti-GAF antibodies to identify protein-specific chromatin interactions at a 1kb resolution. **(B–C) Frequency of indirect looping versus direct looping.** Pie charts showing the proportion of downregulated **(B)** CLAMP-dependent and **(C)** GAF-dependent loops that are a direct or indirect effect of their respective protein knockdown. **(D) Overlap of direct and indirect CLAMP- and GAF-dependent loops.** UpSet plot showing the intersection between directly and indirectly mediated loop categories for both CLAMP and GAF. **(E) Direct loop size distribution shows size specificity over HAS.** Boxplots comparing the genomic span (kb) of indirect (left) and direct (right) loops across High Affinity Sites (HAS), the X chromosome (chrX), and autosomes. Direct loops at HAS exhibit a more constrained size distribution compared to those on autosomes. Asterisks (*) indicate significant difference between direct and indirect loop sets (Mann-Whitney Test, *P* < 0.05). **(F) *De novo* motif enrichment at direct anchors.** Tables listing the top *de novo* motifs at direct CLAMP and GAF loop anchors. **(G) Hi-ChIP Aggregate Peak Analysis (APA).** APA plots showing enrichment of CLAMP and GAF binding at HAS and GA-repeats on the X chromosomes and autosomes relative to random GA-repeats (control). Pixel resolution is 1.0kb with ± 10kb flanking central pixel. **(H) Visualization of direct vs indirect looping on the X chromosome.** CoolBox plots displaying: **(I & III)** Differential contact matrices (log_2_FC) showing changes in TAD architecture following *clamp* and *gaf* RNAi treatment. **(II & IV)** Normalized Hi-ChIP occupancy tracks coupled with annotated direct (blue) and indirect (green) chromatin loops. **(V)** *dm6* gene annotations on the X chromosome.

By integrating Micro-C with Hi-ChIP, we were able to assign with high confidence which downregulated CLAMP-and GAF-dependent loops were directly or indirectly regulated by each factor. Furthermore, we defined direct looping as either factor being present at one or both of the two loop anchors and indirect looping as neither factor being present at either loop anchor. We characterized the following four groups of loops (Fig. 3B, C): 1) direct downregulated CLAMP-dependent loops (N=1,366, 8.3%), 2) indirect downregulated CLAMP-dependent loops (N=15,150, 91.7%), 3) direct downregulated GAF-dependent loops (N=1,679, 30.3%), and 4) indirect downregulated GAF-dependent loops (N=3,870, 69.7%). As we had expected, there are fewer direct than indirect loops consistent with CLAMP and GAF being essential proteins which regulate the expression and function of many other regulators in a cellular context (Urban et al. 2017b; Adkins et al. 2006).

Next, we wanted to know whether there was overlap between the direct and indirect loop sets. We performed an intersection of all loops and determined that, although minimal, there was significant overlap between direct CLAMP- and GAF-dependent loops (N=7, 0.23%) and indirect CLAMP- and GAF-dependent loops (N=39, 0.21%) (Fig. 3D; *P* < 0.001, Fisher’s Exact Test). This indicates that there are a few loci which CLAMP and GAF directly or indirectly influence together. However, given that the magnitude of the overlap is very small, we suggest that these two pioneer TFs possess distinct genome-wide looping roles although they bind to very similar binding sites (Fig. 3D).

We next asked whether loops directly mediated by CLAMP or GAF have a differential size distribution compared with indirect loops and each other. Interestingly, the distribution of loop sizes for the indirect CLAMP- and GAF-dependent loops are all significantly larger and contain wider interquartile ranges in comparison to the corresponding direct loop categories (Fig. 3E; *P* < 0.001, Mann-Whitney Test). Interestingly, we observed that unlike other loop classes direct GAF-dependent loops that overlap HAS have a large and wide interquartile range, similar to the indirect loop set (Fig. 3E, *P* < 0.001, Mann-Whitney Test). This indicates that GAF is directly responsible for a wide range of short- to long-range interactions at the HAS, while promoting more uniform short-to medium-sized interactions at other sites on the X chromosome and autosomes. In contrast, the direct CLAMP-dependent loops exhibit a narrow interquartile range and are short in size, suggesting the direct role of CLAMP is mediating short-range local chromatin loops at the HAS, X chromosome, and autosomes. Lastly, the direct GAF-dependent loops are significantly larger than the direct CLAMP-dependent loops on the X chromosome and autosomes (Fig. 3E; *P* < 0.001, Mann-Whitney Test). Notably, the direct GAF-dependent loops exhibited a larger loop-size range at the HAS than direct CLAMP-dependent loops (Fig. 3E), but the total number of chromatin loops exhibiting this difference was not enough to provide statistical power to the comparison of the medians. Together, these data suggest that CLAMP regulates the formation of a shorter range and largely mutually exclusive set of chromatin loops compared with GAF.

To identify DNA motifs present at directly regulated loop anchors, we intersected CLAMP and GAF ChIP peaks with their respective loop anchors, thereby retaining only TF-bound loop anchors. Subsequently, we utilized the HOMER motif discovery and analysis suite (Heinz et al. 2010) to identify *de novo* motifs. We report the top *de novo* motif found in each site category using the direct CLAMP- and GAF-dependent TF-bound anchors (Fig. 3F). The most striking difference in motifs is between CLAMP- and GAF-dependent anchors at the HAS, where GA-repeats are interrupted by a single highly conserved cytosine nucleotide (67.3% frequency in Motif PWM) on CLAMP-bound anchors but not GAF-bound anchors (Fig. 3F, upper). This single cytosine interruption is also maintained in the top motif detected in CLAMP-dependent anchors on other sites on the X chromosome (61.6% frequency in Motif PWM). In the remaining site categories, the break within the GA-repeat motif is less pronounced as we observed cytosine conservation increase only modestly at other locations in the genome. These results are consistent with our prior findings that a cytosine residue within the GA-repeat favors CLAMP compared with GAF binding (Kaye et al. 2018). Together, these results suggest that CLAMP and GAF bind GA-repeats that differ by a single nucleotide where they drive the formation of loops of different sizes.

Next, we asked whether the pattern of chromatin looping at GA-repeat anchors differs between direct CLAMP-and GAF-dependent loops at HAS and other GA-repeats throughout the genome. Given that GA-repeats are abundant in the genome and have multiple functions, we restricted our analysis to GA-repeats located at the loop anchors of CLAMP or GAF direct loops (either at HAS or other genomic locations), thereby isolating GA-regulatory elements engaged in TF-mediated chromatin looping. Leveraging CLAMP- and GAF-specific Hi-ChIP contact matrices, we used APA plots to measure TF-specific aggregated looping over three classes of sites: HAS, GA-repeats on the X chromosome, and GA-repeats on the autosomes (Fig. 3G). Subsequently, we created union sets of the GA-repeats which interact with CLAMP or GAF since either TF may interact with the HAS.

We observed that CLAMP has a strong enrichment of direct loops at the HAS, with the central pixel (1kb bin) showing substantial contact frequency relative to flanking regions (Fig. 3G). This looping signal is consistent with prior work demonstrating that CLAMP binds directly to the HAS (Soruco et al. 2013; Eggers and Becker 2021), and extends these findings by revealing that CLAMP not only recognizes the HAS but also mediates chromatin looping at these sites. In contrast, GAF showed a broader, less focused looping pattern at the HAS, with enrichment dispersed across a ±3kb window rather than concentrated at the central pixel (Fig. 3G). This diffuse signal, while still centered on the HAS, is consistent with prior reports that GAF contributes to DCC binding but it is not nearly as strongly required as CLAMP (Soruco et al. 2013) (Kaye et al. 2018). Together, these results suggest that the ability of CLAMP to mediate precise, focal chromatin loops at the HAS may be critical for efficient DCC recruitment, whereas GAF may stabilize or reinforce these interactions over a broader spatial range.

Next, we examined direct looping by CLAMP and GAF at GA-repeats on the X chromosome and autosomes (outside of HAS) to define context-specific functions of these factors. CLAMP preferentially regulated looping at X-linked GA-repeats compared to autosomal GA-repeats, consistent with its known enrichment on the male X chromosome (Soruco et al. 2013; Kaye et al. 2018). In contrast, GAF exhibited similar looping strength at GA-repeats on both the X chromosome and autosomes, with only a modest increase on the X, suggesting that GAF functions more uniformly across chromosomes at these elements. Interestingly, aggregated loops directly regulated by CLAMP displayed a striped signal pattern at GA-repeats on both the X chromosome and autosomes (Fig. 3G), indicating that CLAMP promotes broad, directional chromatin interactions stemming from GA-rich loci rather than forming discrete point-to-point loops (Vian et al. 2018). In contrast, loops directly regulated by GAF showed no detectable striping signal at GA-repeats on either chromosome, suggesting that GAF organizes more focal chromatin contacts without the extended directional interactions that CLAMP exhibits. Together, our results suggest that CLAMP and GAF have distinct context-specific direct functions in mediating looping at HAS, X-chromosome, and autosomal GA-repeats.

To visualize the differential characteristics of direct versus indirect CLAMP- and GAF-mediated looping, we examined a 988 kb region of the X chromosome (chrX:3,340,000-4,328,000) containing HAS, TF-specific loops, and differential TADs (Fig. 3H). CLAMP-dependent loops in this region recapitulated global patterns (Fig. 3H: II): direct loops (blue) comprised only a small fraction of total loops and were significantly smaller than indirect loops (green), consistent with genome-wide observations (Fig. 3A). In contrast, GAF-dependent loops exhibited distinct characteristics (Fig. 3H: IV): direct GAF loops showed greater size diversity and more engagement with diverse loci, mirroring the broader size distribution observed at HAS (Fig. 3D, right). Notably, TF occupancy did not always predict direct looping: 66.3% of CLAMP ChIP-seq peaks (N=8,305/12,518) and 55.4% of GAF peaks (N=6,465/11,680) were not associated with Hi-ChIP loops (Fig. 3H: II, IV). These findings indicate that CLAMP and GAF occupancy and direct looping are not always coupled, potentially due to redundant architectural proteins that maintain loops following TF depletion and/or additional functions of CLAMP and GAF beyond chromatin looping.

Notably, the disruption of CLAMP- and GAF-dependent looping resulted in differential TAD formation under both *clamp* and *gaf* RNAi knockdown conditions (Fig. 3H: I, III; Fig. S3). Given that HAS (Fig. 3H, orange highlighted loci) are known to populate TAD boundaries (Ramírez et al. 2015), these findings reinforce the role of these factors in establishing and maintaining TAD integrity. To quantify the relationship between loop loss and TAD reorganization, we compared the differential TADs (Fig. 3H: I, III) with our direct loop sets (Fig. 3H: II, IV), determining what proportion of TAD structural changes could be directly attributed to the loss of CLAMP or GAF loops. Direct CLAMP-dependent loops were associated with 41.7% (N=508/1218) of differential CLAMP TADs, while direct GAF-dependent loops were associated with 32.4% (N=1105/3415) of differential GAF TADs. This substantial overlap between direct loop loss and TAD reorganization provides quantitative evidence that CLAMP and GAF play direct, mechanistic roles in maintaining TAD architecture, rather than acting solely through indirect or compensatory mechanisms.

Lastly, direct CLAMP- and GAF-dependent looping affect a variety of genomic elements as seen by the diverse chromatin states and genes enriched within a specific region (Fig. 3H: V, VI). Therefore, we hypothesized that it may be that these distinct, direct looping contacts regulate interactions between different chromatin states on the X chromosome compared with autosomes.

### 4. GAF regulates chromatin looping at silent euchromatin specifically on the X chromosome and not on autosomes while CLAMP regulates chromatin looping between active chromatin regions throughout the genome

To investigate the distinct functions of CLAMP and GAF in directly regulating chromatin looping on the X chromosome and autosomes, we analyzed sequence elements and chromatin states that are present at directly regulated loop anchors (Fig 4). Interestingly, we observed that while GAF-mediated loops on autosomes predominantly connect active chromatin states (States 1 and 3), GAF predominantly regulates looping at inactive silent euchromatin on the X chromosome (State 9) (Fig. 4A, B). Our data suggest that the function of the GAF-CLAMP competition (Kaye et al. 2018) may be to alter the location of chromatin loops directly regulated by GAF specifically on the X chromosome which promotes CLAMP and not GAF occupancy at active chromatin regions that require dosage compensation (Fig. 3E-H).

**Figure 4.**
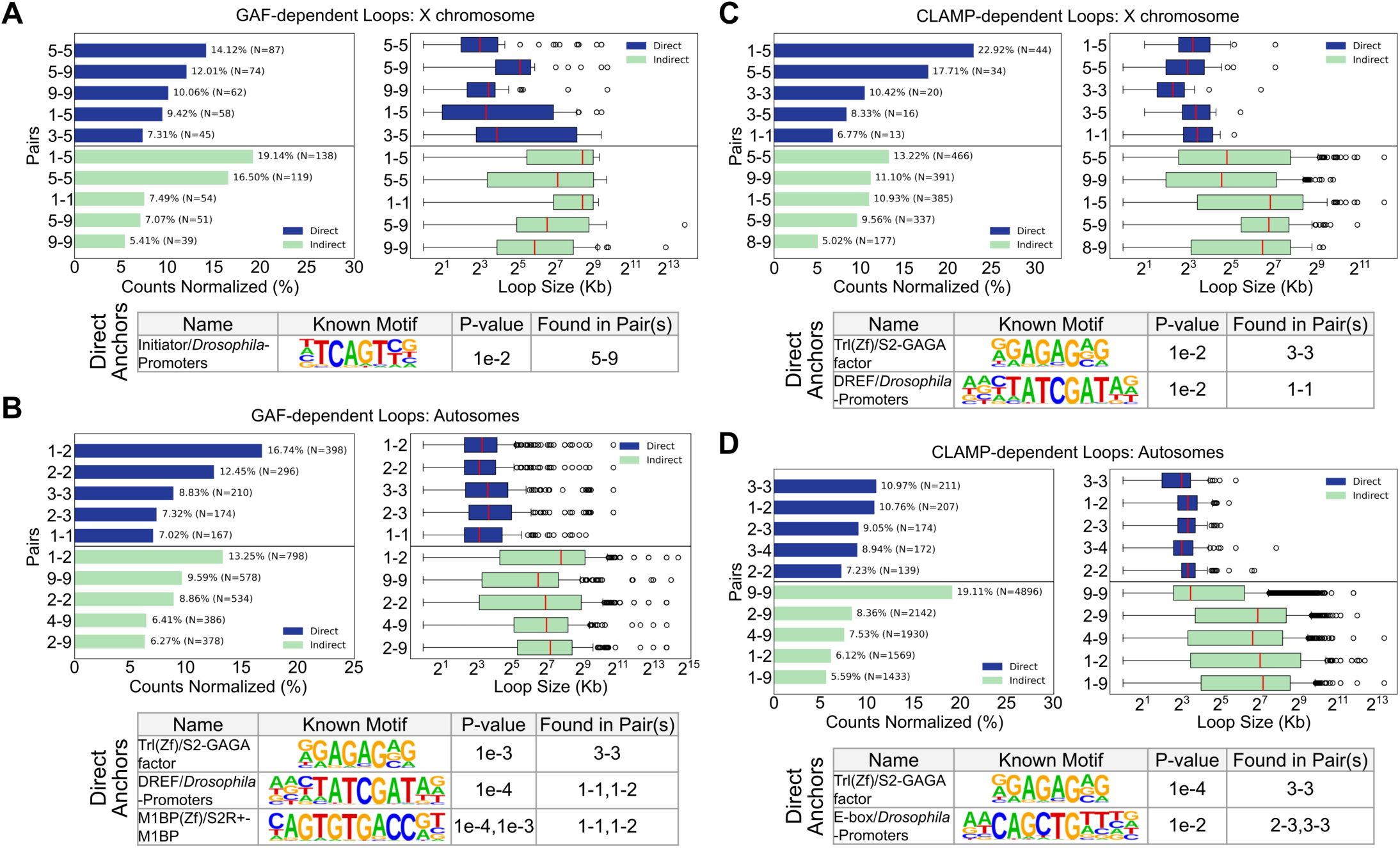
Chromatin State Pair enrichment of CLAMP- and GAF-dependent loops shows GAF regulates chromatin looping at silent euchromatin on the X chromosome and not on autosomes. CLAMP on the (A) X chromosome and (B) autosomes: State Pair Enrichment: Normalized counts (%) of direct (blue) and indirect (green) downregulated loops, categorized by the chromatin state pairs at their anchors. **Loop Size (Top):** Comparison of genomic span (kb) for direct and indirect loops across enriched state pairs. On the **X chromosome**, direct loops frequently bridge State 1 and 3 (Active Promoters and Enhancers) with State 5 (DC genes). **Size Distribution:** Direct interactions consistently exhibit a smaller, more constrained size distribution compared to the larger span of indirect contacts. **Motif Enrichment (Bottom):** Known motif discovery analysis for loop anchor pairs. Significant motifs include the GAGA-factor (Trl) and DREF promoter elements specifically enriched at direct CLAMP-dependent anchors. **(C-D) GAF-mediated chromatin looping. GAF on the (C) X chromosome and (D) autosomes: State Pair Enrichment:** Distribution of direct and indirect GAF-dependent loops across chromatin state pairs. GAF preferentially organizes loops between State 9 (transcriptionally silent euchromatin) and State 5 (DC genes) on the X chromosome, while facilitating active chromatin contacts (States 1-3) on autosomes. **Loop Size (Top):** Comparison of genomic span (kb) for GAF-mediated interactions. Direct GAF loops (blue) maintain a larger genomic span on the X chromosome than on autosomes and indirect GAF loops (green) maintain a broader genomic span across all categories. **Motif Enrichment (Bottom):** Known motif discovery analysis for loop anchor pairs at direct GAF-mediated anchors shows enrichment of Initiator (Inr) motif on the X chromosome while GAGA-factor (Trl) and DREF motifs were enriched on autosomes.

While GAF regulates the formation of chromatin loops between active chromatin states rather than silent regions on autosomes, on the X chromosome, we observed GAF often tethers active transcriptional chromatin states (States 1, 3, and 5) to transcriptionally silent states (State 9) (Fig. 4A, B). Notably, the loop anchors at silent states on the X chromosome, where chromatin looping is directly regulated by GAF, are located within transcriptionally silent intergenic euchromatin and insulator-rich regions (State 9) rather than Polycomb-repressed (State 6) or constitutive heterochromatin (States 7 and 8) (modENCODE Consortium et al. 2010). Therefore, GAF regulates looping that brings DC genes near insulators but not heterochromatin (Fig. 4A, B). We also found that direct GAF-mediated loops have a larger median loop size on the X chromosome (8kb–35kb), where GAF is outcompeted by CLAMP, than on autosomes (9-13kb) (Fig. 4A, B) (Kaye et al. 2018). Overall, chromatin loops that are regulated by GAF on the X chromosome differ significantly from those on autosomes because they are enriched for direct links between insulators located within silent euchromatin and other genomic loci (Fig. 4A, B).

In contrast to GAF, CLAMP directly mediates chromatin looping between distal enhancers (State 3) and promoters (State 1) and DC genes on the X chromosome (State 5) (Fig. 4C). On autosomes, CLAMP direct loops also bridge active chromatin states (Fig. 4D). The median loop size of direct CLAMP loops is 8-11kb on the X chromosome and 8-10kb on autosomes, suggesting that CLAMP regulates loops of similar sizes on the X chromosome and autosomes (Fig. 4C, D). Therefore, CLAMP directly facilitates precise, short-range chromatin interactions between transcriptionally active chromatin and dosage-compensated genes on the X chromosome and between active genes on autosomes.

Distinguishing the direct from indirect effects of GAF and CLAMP is critical because direct and indirect chromatin loops occur at very different chromatin states. While the chromatin loops that are directly facilitated by CLAMP and GAF are distinct from each other in scale and chromatin state, the majority of chromatin loops that change after the loss of either factor are indirectly regulated (Fig. 3B, C). Furthermore, there are differences between the loops that are indirectly regulated by CLAMP and GAF: we observed that while CLAMP directly mediates chromatin loops *only* between loci within active states (States 1, 3, and 5), loops indirectly regulated by CLAMP include inactive states (States 8 and 9) (Fig. 4C). Conversely, loops indirectly regulated by GAF include an active state (State 1) on the X chromosome which was not observed in the directly regulated chromatin loops (Fig. 4A). Therefore, it is key to distinguish direct from indirect effects to understand the differential functions of CLAMP and GAF. Future analysis will define how the loss of CLAMP or GAF indirectly regulates chromatin looping through modulating the function of other looping factors.

To identify potential cofactors that may function with CLAMP or GAF, we performed HOMER motif analysis on direct CLAMP and GAF loop anchors genome-wide. As expected, we find the Trl(Zf)/GAGA-factor motif (*P* = 1e-4; Fig. 4D) at CLAMP-mediated 3-3 loop anchors on X (*P* = 1e-2; Fig. 4C) and autosomes (*P* = 1e-4; Fig. 4D), likely due to the high sequence similarity between CLAMP and GAF (Fig. 3F). Additionally, at CLAMP-mediated loops on the X chromosome, we found the DNA replication-related element factor (DREF) binding motif (Fig. 4C), a factor which binds and activates housekeeping genes (Zabidi et al. 2015). The enrichment of the DREF motif at promoter-promoter (1-1) contacts (*P* = 1e-2; Fig. 4C) suggest that loop anchors that are directly regulated by CLAMP are enriched for housekeeping genes on the X chromosome, which are the first genes to be compensated during development (Prayitno et al. 2019). In contrast, on autosomes, CLAMP regulates chromatin loop anchors enriched for E-box enhancer elements at enhancer–enhancer (3-3) and exon–enhancer (2-3) contacts (*P* = 1e-2; Fig. 4D), where developmental genes would require canonical E-box on-off switches (Gordân et al. 2013).

We next examined motifs associated with chromatin loops directly regulated by GAF. GAF-mediated loops on the X chromosome that connect dosage compensation genes (state 5) with silent euchromatin insulator regions (State 9) are enriched for the Initiator (Inr) motif (Fig 4A). Inr is a promoter element that often cooperates other core promoter elements as it cannot initiate transcription alone (Vo Ngoc et al. 2019). It is possible that at proximal promoter regions of DC genes, GAF may require additional factors, such as co-promoter element Inr, to facilitate looping on the X chromosome.

Unlike the X-chromosome loop anchors, autosomal GAF-mediated loop anchors are enriched for motifs exclusively within active chromatin states (States 1-3) (Fig. 4B). The presence of DREF and Motif-1-binding protein (M1BP), an RNA Pol II pausing factor associated with housekeeping genes (Poliacikova et al. 2023), at active promoter–exon (1-3) contacts suggests a distinct autosomal role for GAF (Fig. 4B). On autosomes, GAF may facilitate looping to regulate housekeeping genes consistent with prior work (Li and Gilmour 2013). Overall, CLAMP direct loop anchors are at housekeeping genes on the X chromosome that need to be dosage-compensated, and GAF direct loop anchors are at housekeeping genes on autosomes suggesting that the CLAMP-GAF competition may allow CLAMP to be enriched at X-linked housekeeping genes instead of GAF to allow these genes to recruit the DCC.

We next compared the size of different classes of loops connecting different chromatin states on the X chromosome and autosomes (Fig. 4). We observed that on the X chromosome, GAF-mediated state 5 (dosage compensation)-state 9 (insulators) loops are significantly larger than state 5-5 (*P* = 3.83e-8) or state 9-9 (*P* = 1.94e-8) loops (Fig. 4A). Additionally, state 9-9 loops were significantly smaller than state 3-5 loops (*P* = 1.84e-2; Fig. 4A). Therefore, it is possible that GAF mediates larger state 5-9 chromatin loops that connect insulators to dosage compensated genes to provide a three-dimensional scaffold for the dosage compensation molecular hub. This larger GAF-mediated loop scaffold may explain the function of GAF in dosage compensation that is weaker than that of CLAMP but still important for DCC recruitment (Kaye et al. 2018).

Collectively, our results reveal that CLAMP and GAF deploy context-dependent strategies to organize chromatin architecture: CLAMP directly mediates short-range loops between active chromatin states on both X and autosomes. In contrast, the function of GAF diverges based on chromosomal context, facilitating compact loops between active loci on autosomes, but forming larger loops between dosage-compensated genes and silent euchromatin on the X chromosome. These distinct looping patterns reflect the competitive dynamics between CLAMP and GAF on the X chromosome, where the exclusion of GAF from active sites by CLAMP likely redirects it to silent regions where it regulates larger loops that may provide structural scaffolding for DCC spreading. Our findings demonstrate how competitive binding enables CLAMP and GAF to context-specifically regulate chromatin loops that are distinct in both size and chromatin state.

### 5. Chromatin loops directly mediated by CLAMP and GAF associate with *roX2* chromatin loops on the X chromosome

The DCC incorporates one of two long non-coding RNAs (lncRNAs), *roX1* or *roX2*, and therefore these lncRNAs are spatial markers of DCC localization (Meller and Rattner 2002). Tian et al. (2024) performed the RNA-associated chromatin DNA-DNA interaction (RDD) method targeting *roX2* across the genome in *Drosophila* S2 cells to capture chromatin loops associated with this lncRNA and observed the expected X-chromosome specificity. Furthermore, we recently demonstrated that CLAMP can directly interact with *roX2 in vivo* and *in vitro* (Ray, M*, Zaborowsky, J*, Mahableshwarkar, P, Aguilera, J, Huang, A, Vaidyanathan, S, Fawzi, N and Larschan, E. 2024). We therefore leveraged these *roX2*-associated loops as a proxy for three-dimensional DCC localization and spreading across the X chromosome, enabling us to determine the proportion of direct CLAMP- and GAF-dependent loops that spatially associate with the DCC beyond the HAS. Prior to integrating our differential loop datasets with the *roX2*-RDD chromatin loops, we equalized bin sizes across all datasets to enable direct comparison (see Methods), and we report the complete loop counts after normalization (Table S2).

We first asked what proportion of *roX2*-RDD loops on the X chromosome overlap with direct CLAMP- and GAF-dependent loops, and whether this proportion is enriched compared to indirect loops. For CLAMP-dependent loops, direct loops overlapped with *roX2*-RDD loops significantly more than their corresponding indirect loops at HAS (5.2% vs 2.0%; *P* < 0.05, Fisher’s Exact Test) and on the X chromosome (20.7% vs 5.5%; *P* < 0.05, Fisher’s Exact Test) (Fig. 5A). The same pattern was observed for GAF-dependent loops, with direct loops similarly enriched compared with indirect loops at HAS (3.7% vs 1.5%; P < 0.05, Fisher’s Exact Test) and on the X chromosome (12.3% vs 4.2%; P < 0.05, Fisher’s Exact Test) (Fig. 5A). These results suggest that under normal conditions where dosage compensation is active, *roX2*-RDD loops at HAS and on the X chromosome spatially associate with direct CLAMP- and GAF-dependent regulation, linking these factors spatially to the DCC. However, the majority of loops in each dataset do not overlap which is consistent with *roX2* spreading to active gene bodies where CLAMP and GAF largely do not localize, and the well-characterized functions of both TFs beyond dosage compensation.

**Figure 5.**
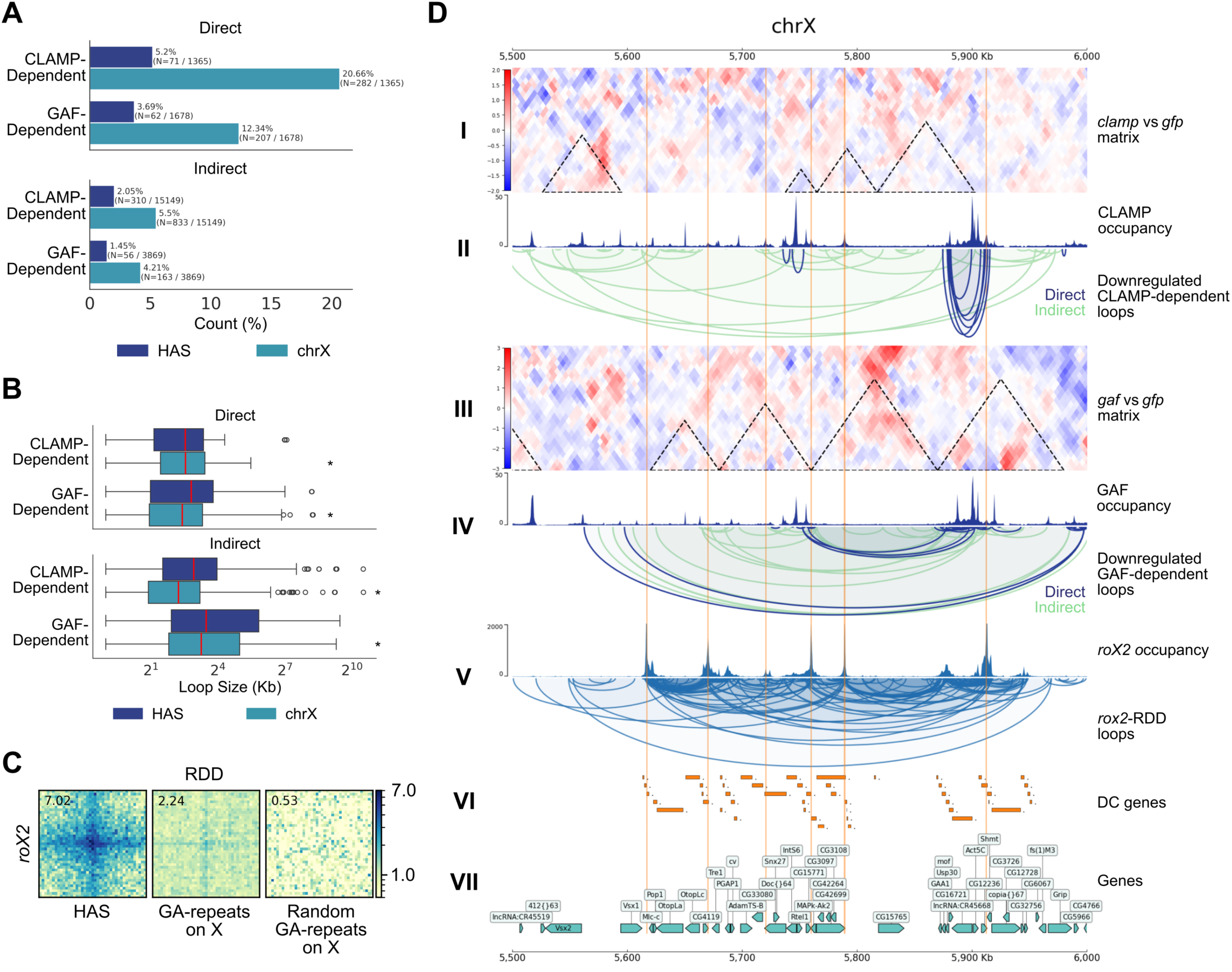
Direct CLAMP- and GAF-dependent loops are associated with a subset of *roX2* chromatin loops on the X chromosome. **(A) Distribution of direct and indirect loops associated with *roX2* loops at HAS and across the X chromosome.** Bar plots showing the normalized counts (%) of direct and indirect CLAMP- and GAF-dependent loops that overlap with *roX2* loops at High Affinity Sites (HAS; dark blue) compared to the X chromosome (chrX; cyan). **(B) Loop size of *roX2*-associated loops.** Box plots comparing the loop size (kb) for direct and indirect CLAMP- and GAF-dependent interactions at HAS versus chrX. **(C) Aggregate Peak Analysis (APA) of *roX2* loops over HAS.** APA plots visualizing *roX2* RNA-associated chromatin DNA-DNA (RDD) interactions at HAS, GA-repeats on the X, and random GA-repeats on the X. Strong enrichment at HAS (7.02) shows that *roX2* binds preferentially at HAS. Pixel resolution is 1.0kb with ± 20kb flanking central pixel. **(D) Co-localization of CLAMP- and GAF-dependent loops with *roX2*-RDD loops.** Representative CoolBox plots of a 500kb genomic window shows **(I & III)** differential contact matrices (log_2_FC) for *clamp* RNAi and *gaf* RNAi versus *gfp* RNAi control, respectively. **(II & IV)** Hi-ChIP CLAMP and GAF protein occupancy tracks aligned with downregulated direct and indirect chromatin loops. **(V)** *roX2* occupancy aligned with *roX2*-RDD loops. **(VI)** Genomic tracks for Dosage Compensation (DC) genes and **(VII)** *dm6* gene annotations on the X chromosome.

Next, given our earlier observation of TF-specific loop size distributions (Figs. 3, 4), we asked whether the *roX2*-associated direct loops exhibit distinct size characteristics compared to indirect loops. On the X chromosome, the loop size distributions of direct and indirect sets were significantly different for both CLAMP- and GAF-dependent loops (*P* < 0.05, Mann-Whitney Test), though this difference was not significant at HAS (Fig. 5B). Specifically, direct CLAMP- and GAF-dependent loops both exhibited a narrower range of loop sizes than indirect loops (Fig. 5B), consistent with the local looping patterns observed previously (Fig. 3). The preferential overlap of direct CLAMP- and GAF-dependent loops with *roX2*-RDD loops, combined with their shorter-range size profiles and reduced overlap at HAS, suggests that these factors may contribute to the local spreading of dosage compensation on the X chromosome.

Having established the unique CLAMP- and GAF-specific looping characteristics at HAS and GA-repeats (Fig. 3), we next asked how *roX2*-RDD loops aggregate over these same classes of sites on the X chromosome (Fig. 5C). Using APA plots, we observed strong aggregated *roX2*-RDD looping at HAS, recapitulating the central role of this lncRNA and the DCC in X-chromosome dosage compensation (Fig. 5C). We then examined *roX2*-RDD looping over GA-repeats that directly associate with CLAMP and GAF on the X chromosome. Notably, aggregated *roX2*-RDD looping and direct CLAMP-dependent looping at both HAS and across the X chromosome exhibited a similar striped signal pattern, indicating that *roX2*-RDD loops, like direct CLAMP-dependent loops, favor broad, directional chromatin interactions emanating from GA-rich loci rather than forming discrete point-to-point contacts (Fig. 3F, 5C). However, aggregated *roX2*-RDD looping at HAS (log enrichment = 7.02) was substantially greater than that of either direct CLAMP- or GAF-dependent loops (log enrichment = 1.80 and 1.68 respectively) (Fig. 5C). These results suggest that HAS serve as key regulatory hubs for dosage compensation, and that *roX2*-associated looping more closely resembles direct CLAMP-dependent looping than GAF-dependent looping, consistent with our previous work demonstrating how the DCC (which includes *roX2)* and CLAMP may cooperate at HAS to mediate the local spreading of dosage compensation on the X chromosome (Ray, M*, Zaborowsky, J*, Mahableshwarkar, P, Aguilera, J, Huang, A, Vaidyanathan, S, Fawzi, N and Larschan, E. 2024; Soruco et al. 2013).

Lastly, to visualize *roX2*-RDD loops alongside direct CLAMP- and GAF-dependent loops, we examined a representative region on the X chromosome (chrX:5,500,000-6,000,000) (Fig. 5D). Within this region, differential CLAMP- and GAF-dependent loops exhibited looping characteristics consistent with genome-wide patterns (Fig. 3D, Fig. 5D: II, IV). The *roX2*-RDD loops encompassed a broad size distribution at this locus (Fig. 5D: V), a pattern also observed genome-wide (Fig. S4), spanning both intra- and inter-TAD interactions (Fig. 5D: I, III, V). Notably, *roX2* and CLAMP occupancy were enriched at HAS (orange highlighted loci), while GAF occupancy was not (Fig. 5D: II, IV, V), further supporting the closer functional relationship between *roX2* and CLAMP at these sites compared with GAF. In addition, *roX2*-RDD loops were present at every control TAD on the X chromosome, suggesting that *roX2*-associated looping is a ubiquitous feature of X-chromosome architecture essential for dosage compensation. These *roX2*-RDD loops also encompassed all genes undergoing dosage compensation (Fig. S5, Fig. 5D: VI, VII), consistent with a direct spatial relationship between *roX2*-associated chromatin architecture and the transcriptional output of the DCC. Together, we demonstrate that *roX2*-RDD loops span the X chromosome and spatially associate with direct CLAMP- and GAF-dependent loops, illustrating how these factor-specific loops shape the three-dimensional context for dosage compensation.

## Discussion

A fundamental question about gene regulation across species is: How are the correct genes targeted for coordinate regulation within the highly compact three-dimensional nucleus? We used the upregulation of X-linked genes (XCU) as a model to address this question because we have identified key DNA sequences (Alekseyenko et al. 2008, 2012) and transcription factors (TFs) (Soruco et al. 2013; Larschan et al. 2012) that are involved in targeting the DCC specifically to the X-chromosome, including competition between two essential pioneer TFs (Kaye et al. 2018). However, the mechanisms by which sequence elements and TFs function directly to establish an X-specific three-dimensional chromatin environment remained unknown. Here, we provide mechanistic evidence to support the following model for X-chromosome specificity (Figure 6): 1) On the X-chromosome, a single base change within GA-rich TF binding sites alters the relative affinity of CLAMP and GAF which then favors the formation of smaller CLAMP driven loops at active dosage-compensated housekeeping genes and larger GAF-driven loops at silent insulator regions. 2) In contrast, on autosomes both CLAMP and GAF mediate loops between active genes, although GAF-regulated loops have a broader range of sizes than CLAMP-regulated loops. Overall, we provide insight into how the X chromosome is recognized for dosage compensation.

**Figure 6.**
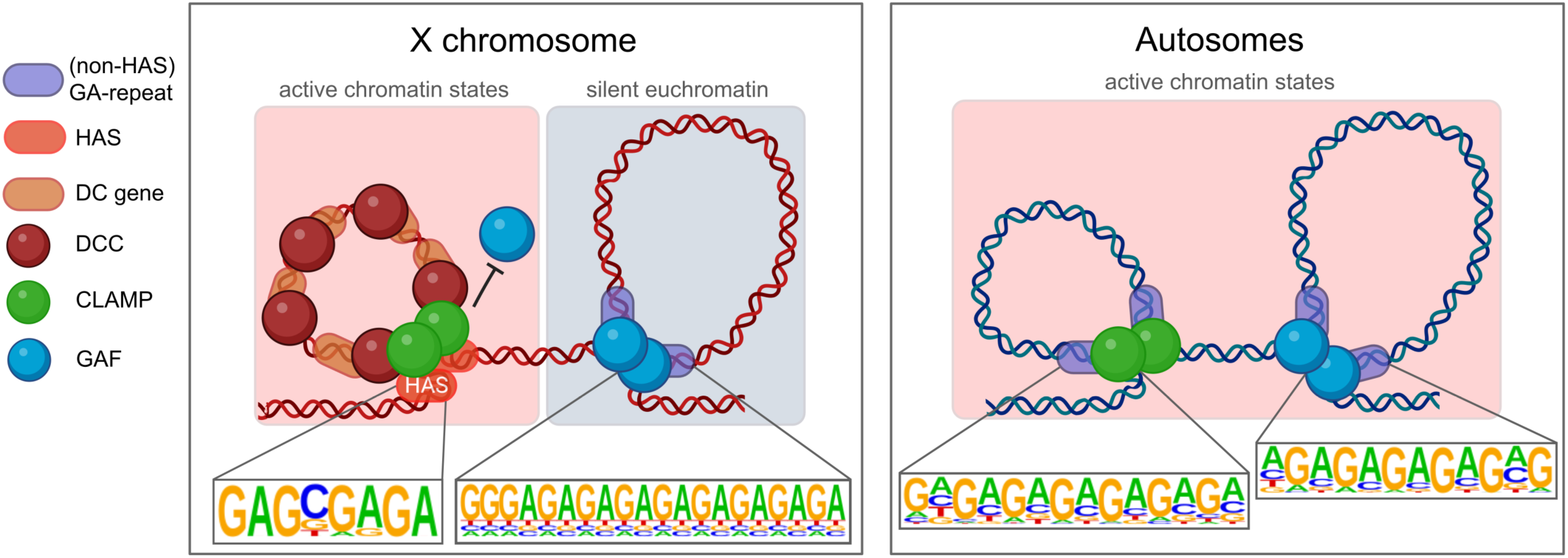
Model for X-specific CLAMP and GAF function in chromatin looping. **(1)** On the X chromosome, a single base change within GA-rich TF binding sites alters the relative affinity of CLAMP and GAF which then favors the formation of smaller CLAMP driven loops at active dosage-compensated housekeeping genes and larger GAF-driven loops at silent insulator regions. **(2)** In contrast, on autosomes both CLAMP and GAF mediate loops between active genes, although GAF-regulated loops have a broader range of sizes than CLAMP-regulated loops.

There are many contexts in which the DNA sequence content of TF binding sites varies, including within SNPs related to disease (Yan et al. 2021; Khetan et al. 2025). Our data suggest that even single base alterations of TF binding sites can alter three-dimensional nuclear organization by favoring the binding of one TF instead of another which can drive different transcriptional programs. For example, the accumulation of binding sites on the X chromosome that favor CLAMP and not GAF near housekeeping genes that need to be dosage compensated promotes an X-specific three-dimensional organizational hub that allows CLAMP to recruit the DCC to genes that need to be dosage compensated.

GAF also contributes to DCC recruitment but to a much lesser extent than CLAMP (Kaye et al. 2018). Based on our three-dimensional analysis of the direct function of GAF, it is possible that GAF contributes to DCC recruitment by generating an X-specific three-dimensional scaffold consisting of larger chromatin loops between X-linked insulators. In contrast, on autosomes GAF regulates interactions between active chromatin regions, such as enhancers and promoters. This context-specific architectural loop function is consistent with prior reports that suggest that GAF is an important component of multiple insulator complexes (Lomaev et al. 2017; Wolle et al. 2015) and has a function in dosage compensation (Kaye et al. 2018, 2017; Greenberg et al. 2004). Furthermore, GAF also mediates several very long-range looping interactions in the brain that are important for regulating functionally similar genes that are encoded at different locations in the genome (Li et al. 2023). Therefore, our data provide mechanistic insight to explain prior observations that GAF functions as an insulator protein and has a role in dosage compensation.

In contrast to the function of GAF at insulator loci on the X chromosome where it facilitates long-range interactions, we demonstrate that CLAMP facilitates small scale loops between active regions on both the X chromosome and autosomes. It is possible that these smaller hubs are critical for the spreading of the DCC from the HAS to the bodies of active genes where it increases transcript levels. Interestingly, the first genes to be compensated that are in close two-dimensional proximity to the HAS are the housekeeping genes which encode components of essential protein complexes that require precise stoichiometry (Prayitno et al. 2019). Housekeeping genes are often enriched for the DREF motif (Hochheimer et al. 2002) which we found as highly enriched at loops which were directly mediated by CLAMP but not GAF on the X-chromosome and at autosomal loops regulated by GAF and not CLAMP. This inverse relationship between looping at housekeeping genes on the X chromosome and autosomes suggests that competition between CLAMP and GAF is important to assure that CLAMP but not GAF is enriched at X-linked housekeeping genes that require dosage compensation.

There are many possible biomolecular mechanisms by which CLAMP and GAF could regulate mutually exclusive sets of looping interactions. One possibility is that their intrinsically disordered domains (IDRs) that do not bind DNA have an independent set of interactors and biophysical properties. For example, the CLAMP IDR interacts with both the *roX* RNAs (*in vivo* and *in vitro*) (Ray, M*, Zaborowsky, J*, Mahableshwarkar, P, Aguilera, J, Huang, A, Vaidyanathan, S, Fawzi, N and Larschan, E. 2024) and MSL2 (Eggers et al. 2023; Valsecchi et al. 2020; Tikhonova et al. 2019) which are not known GAF interactors. Furthermore, our work is consistent with a very recent study that indicates that heterotypic interactions between a wide variety of sequence motifs including the GA-rich motif are important for boundary formation in *Drosophila (Varisco et al. 2025)*.

The function of loop extrusion in defining three-dimensional organization in *Drosophila* had been debated until a very recent study quantitatively defined the function of loop extrusion compared to that of tethering elements which can be regulated by GAF (Choppakatla et al. 2026). Tethering elements and loop extrusion are both important for chromatin loop formation, but they have different roles. Loop extrusion functions more strongly during the process by which enhancers and promoters are searching for the correct contacts whereas tethering elements function to stabilize these contacts and regulate transcriptional output. Our work suggests that the specific TFs that bind to tethering elements control the size and chromatin context of the loops that are stabilized. Overall, we provide mechanistic insight into how specific genomic loci are targeted for coordinated gene regulation within a highly compacted three-dimensional chromatin environment.

## Methods

### Cell culture

S2 cells were started from frozen stocks obtained from the *Drosophila* Genomics Resource Center and maintained at 25°C in Schneider’s media supplemented with 10% Fetal Bovine Serum and 1.4X Antibiotic-Antimycotic (Thermofisher Scientific, USA). Cells were passaged every 3 days to maintain an appropriate cell density.

### Western Blot

Protein was extracted from cell pellets using Cell Lysis Buffer (Cell Signaling Technology, 9803) supplemented with 1 mM protease inhibitor cocktail (Abcam, ab65621). Protein concentrations were determined using the DC Protein Assay (Bio-Rad, 5000116). Equal amounts of protein were mixed with NuPAGE™ Sample Reducing Agent (Thermo Fisher Scientific, NP009) and NuPAGE™ LDS Sample Buffer (Thermo Fisher Scientific, NP0007), boiled at 70 °C for 10 min, and loaded onto a 4%–12% gradient SurePAGE™ Bis-Tris gel (GenScript, M00652). Proteins were transferred to a methanol-activated PVDF membrane using a semi-dry transfer method with the Trans-Blot Turbo RTA Transfer Kit PVDF (Bio-Rad, 1704273), Trans-Blot Turbo 5× Transfer Buffer (Bio-Rad, 10026938), and the Bio-Rad Trans-Blot Turbo Transfer System (1.3A, 25 V) for 10 min. Membranes were blocked in 5% milk in phosphate-buffered saline containing 0.05% Tween-20 (PBST) for 20 min at room temperature, then incubated with primary antibodies diluted in 5% milk in PBST overnight at 4 °C. Membranes were washed with PBST and incubated with HRP-conjugated secondary antibodies diluted in 5% milk in PBST for 1 h at room temperature, followed by additional washes in PBST. Protein bands were detected using SuperSignal™ West Femto Maximum Sensitivity Substrate (Thermo Fisher Scientific, 34095) and imaged with the Bio-Rad ChemiDoc Imaging System. GAPDH was used as a loading control and was detected using an HRP-conjugated GAPDH monoclonal antibody. All blots are shown in the Supplementary Material. Antibodies and dilutions were as follows: GAPDH (Proteintech, HRP-60004, 1:1000), CLAMP (1:1000), GAF (1:1000), Anti-Rabbit (Cell Signaling, 7074S, 1:1000).

### Micro-C

Micro-C was performed on S2 cells using two biological replicates for each of the following conditions: *clamp* RNAi, *gaf* RNAi, and *gfp* RNAi (control). For each replicate, 3×10^6^ cells were incubated with 15 μg of the corresponding dsRNA, resulting in six samples total. S2 cells were treated in 6-well plates with 15μg of dsRNA (*clamp*, *gaf*, or *gfp* control) in 500μl of water. S2 cells were first incubated with the dsRNA in Schneider’s media without FBS or Antibiotic-Antimycotic for 45 minutes. After 45 minutes, 3 mL of medium supplemented with 10% FBS and Antibiotic-Antimycotic was added to each well. The plates were then incubated at 25°C for six days before harvesting. After incubation, cells in each well were scraped, aliquoted into 1.5 ml tubes, then spun at 500g in a swinging bucket rotor for 5 minutes to pellet the cells. Supernatant was removed and cell pellets were washed with 1x PBS, counted, then 3×10^6^ cells (protocol-recommended cell count of 1×10^6^ was optimized to account for reduced size of *Drosophila* cells) were re-pelleted in 1.5ml tubes and placed in −80°C for 1 hour as per the manufacturer’s protocol (Dovetail™ Micro-C Kit user guide version 2.0). To prepare samples, cells were thawed at RT and resuspended in 200 μL of 1x PBS (ThermoFisher Scientific, 10010023) and 2 μL 0.3 M DSG. 37% formaldehyde was then added after a 10-minute incubation on a tube rotator to complete crosslinking, as per the manufacturer’s protocol (Dovetail™ Micro-C Kit user guide version 2.0). Sample digestion, quantification (Tapestation HS D5000), proximity ligation, library preparation, ligation capture, and amplification were done using the Dovetail™ Micro-C Kit (Catalog # 21006), the Dovetail^TM^ Library Module for Illumina (Catalog # 25004), and the primer set for NGS library preparation (DovetailTM Dual Index Primer Set #1 for Illumina, Catalog #25010). Libraries were sequenced with Illumina 150×2 bp pair-end sequencing at 80X coverage. Data was submitted to Gene Expression Omnibus (<u>GSE322508</u>).

### Hi-ChIP

S2 Cells were allowed to grow to confluence then harvested. Two replicates of 10×10^6^ S2 cells were used per HiChIP reaction. Cells were crosslinked using 0.3M DSG and 37% formaldehyde as per the manufacturer’s protocol (Dovetail™ HiChIP MNase Kit user guide). Digestion, lysate preparation, chromatin immunoprecipitation, proximity ligation, and library preparation was conducted using the Dovetail™ HiChIP MNase Kit (Catalog # 21007) followed by the Dovetail Library Prep Kit and the primer set for NGS library preparation (included in kit). For CLAMP and GAF immunoprecipitation, Rabbit anti-CLAMP (5μg/sample, SDIX) and Rabbit anti-GAF (5μg/sample) were used, respectively. Libraries were subjected to Illumina 150×2 bp pair-end sequencing. Data was submitted to Gene Expression Omnibus (<u>GSE322509</u>).

### Micro-C Preprocessing

Micro-C raw data was analyzed using the HiC-Pro toolkit (v3.1.0) (Servant et al. 2015) with default parameters. Normalized contact matrices deriving from HiC-Pro were converted to the cooler format using utility scripts (https://github.com/nservant/HiC-Pro/tree/master/bin/utils) in the HiC-Pro toolkit (v3.1.0) and replicates were subsequently merged.

### Hi-ChIP Preprocessing

HiChIP raw data was analyzed using the Dovetail genomics Hi-ChIP documentation (https://hichip.readthedocs.io/en/latest/index.html). Sequences were mapped to the dm6 genome using the Burrows-Wheeler Aligner (v0.7.17) (Li and Durbin 2009; Li 2013) passing the following arguments: mem, −5, -S, -P, -T0; default values were used for all other settings. Using the parse module of pairtools (v0.3.0) (Open2C et al. 2024), valid ligation events were recorded; the following parameters were changed from default values: --min-mapq 40, --walks-policy 5unique, --max-inter-align-gap 30. PCR duplicates were removed using the dedup module of pairtools (v0.3.0) (Open2C et al. 2024). Subsequently, the split module of pairtools was used to generate the final bam and pair files. The samtools (v1.17) (Danecek et al. 2021) sort module was used to sort the bam files. To determine the quality of the proximity ligation events, scripts generated by the Dovetail Genomics team were utilized, which are deposited in the following GitHub repository (https://github.com/dovetail-genomics/HiChiP).

The cooler suite (v0.9.3) (Abdennur and Mirny 2020) was used to generate contact maps from pair files and replicates were subsequently merged. To call loops, the flexible FitHiChIP tool (v10.0) (Bhattacharyya et al. 2019) was used as resolutions are easily adjustable, and it can detect different categories of 3D chromatin interactions. Furthermore, since FitHiChIP required 1D peaks from the same tissue and conditions, MACS2 (v2.2.9.1) (Zhang et al. 2008) was used to call peaks on the Hi-ChIP data. Consensus loops were created for each condition and were used throughout the study.

### Computational Analysis

#### a Topologically-associated domain (TAD) calling and differential analysis (Micro-C)

TADs were identified for all Micro-C conditions (*gfp*, *clamp*, and *gaf* RNAi) using HiCExplorer (v3.7.5) (Wolff et al. 2020, 2018; Ramírez et al. 2018). TADs were called at six resolutions (2.5kb, 5kb, 10kb, 20kb, 40kb, and 80kb) using the *hicFindTADs* tool with the following parameters: --minDepth RESOLUTION × 3, --maxDepth RESOLUTION × 10, --step RESOLUTION, and -- correctForMultipleTesting fdr. Domains across all resolutions were then merged using hicMergeDomains with default parameters to generate multi-resolution consensus TAD sets for each RNAi condition.

To assess how TAD structure was altered upon knockdown of CLAMP or GAF, we used the *gfp* RNAi condition as the reference and performed differential TAD analysis using the following HiCExplorer (v3.7.5) (Wolff et al. 2020, 2018; Ramírez et al. 2018) workflow: (1) hicNormalize (--normalize smallest; default parameters) to normalize contact matrices across conditions; (2) hicCorrectMatrix (default parameters) to remove systematic biases via iterative correction (ICE); and (3) hicDifferentialTAD (--targetMatrix TREATMENT_MATRIX, --controlMatrix CONTROL_MATRIX, --tadDomains CONTROL_TADS, --pValue 0.05, --mode all; default parameters) to identify differential TADs relative to the *gfp* RNAi reference. Differential TAD analysis was performed at each of the six resolutions described above, and results were subsequently merged.

#### b Multi-resolution differential chromatin loop analysis (Micro-C)

Merged cooler contact maps that were produced for each condition were used to identify differential chromatin interactions using the HiCcompare library (v1.24.0) (Stansfield et al. 2018) in Bioconductor (v3.18) (Huber et al. 2015). Differential chromatin interactions were determined at three different resolutions: 1kb, 5kb, and 10kb. Consensus differential chromatin interactions were constructed by merging all three resolutions using a custom Python script that followed the *hicMergeLoops* method in HiCExplorer (v3.7.5) (Wolff et al. 2020, 2018; Ramírez et al. 2018). Consensus significant differential chromatin interactions (*P* < 0.01) were selected and used throughout the study.

#### c TAD and differential loop integration (Micro-C)

Differential TADs were integrated with the consensus merged sets of differential chromatin interactions using the *pybedtools* suite (v0.11.0) (Dale et al. 2011; Quinlan and Hall 2010). Chromatin interactions in BEDPE format were overlapped with TADs in BED format using the *pairToBed* tool with the following arguments: -type either (default parameters), such that a TAD was considered associated with a given interaction if either anchor of the interaction pair overlapped the TAD. In this manner, each set of differential chromatin interactions was assigned its associated TADs, enabling downstream characterization of TAD-level changes linked to specific loop alterations.

#### d High-affinity site integration (Micro-C)

High-affinity sites (HAS) were integrated with the consensus merged sets of differential chromatin interactions using the *pybedtools* suite (v0.11.0) (Dale et al. 2011; Quinlan and Hall 2010). Chromatin interactions in BEDPE format were overlapped with HAS in BED format using the *pairToBed* tool with the following arguments: -type either (default parameters), such that a HAS was considered associated with a given interaction if either anchor of the interaction pair overlapped the HAS. In this manner, each set of differential chromatin interactions was assigned its associated HAS, enabling downstream characterization of HAS-level changes linked to specific loop alterations.

#### e Dosage-compensated gene integration (Micro-C)

To identify dosage-compensated genes (DCGs) in the *Drosophila* genome, we used the *Drosophila* S2 chromatin 9-state model (Kharchenko et al. 2010). Specifically, state 5, which corresponds to actively transcribed exons on the male X chromosome undergoing dosage compensation, was used to annotate genes subject to dosage compensation. Using the *pybedtools* suite (v0.11.0) (Dale et al. 2011; Quinlan and Hall 2010), we overlapped the chromatin state annotations with *Drosophila* gene coordinates using the intersect tool with the -wa and -wb parameters to retain both overlapping regions. All genes overlapping with state 5 were extracted, deduplicated, and designated as DCGs.

DCGs were then integrated with the consensus merged sets of differential chromatin interactions using *pybedtools*. Chromatin interactions in BEDPE format were overlapped with DCGs in BED format using the *pairToBed* tool with-type either (default parameters), such that a DCG was considered associated with a given interaction if either anchor of the interaction pair overlapped the gene. Additionally, we performed a second overlap using -type ospan (outer span), which captures genes falling entirely within the genomic span of an interaction rather than overlapping an anchor. This allowed us to differentiate between DCGs contacted by loop anchors and those encompassed within loops on the X chromosome. Together, these approaches enabled comprehensive characterization of DCG-level changes linked to specific loop alterations.

#### f Differential Micro-C chromatin loop annotation using Hi-ChIP chromatin loops (Micro-C, Hi-ChIP)

Differential Micro-C chromatin interactions were annotated using transcription factor (TF)-specific Hi-ChIP chromatin interactions for CLAMP and GAF. Because the Hi-ChIP experiments were performed exclusively under control conditions, only the downregulated Micro-C chromatin interactions in each knockdown condition were annotated. Each set of downregulated Micro-C interactions was overlapped with the corresponding Hi-ChIP dataset using the *pairToPair* tool (-type both; default parameters) within the *pybedtools* suite (v0.11.0) (Dale et al. 2011; Quinlan and Hall 2010). The Hi-ChIP interaction sets were generated at 1kb resolution, whereas the Micro-C differential interaction sets spanned 1 kb, 5kb, and 10kb resolutions. To standardize the comparison, we extended the overlap windows for the 1kb and 5kb Micro-C interaction sets using -slop 4500 and -slop 2500, respectively, thereby enabling all overlaps to be performed using uniform 10 kb windows. Downregulated Micro-C interactions that overlapped with Hi-ChIP interactions were classified as TF-specific direct interactions, while those without overlap were classified as indirect interactions, which are interactions lost as a downstream consequence of TF knockdown rather than through direct TF-mediated contact.

#### g Aggregated peak analysis plots (Micro-C, Hi-ChIP)

Micro-C differential chromatin interactions and Hi-ChIP TF-specific chromatin interactions were visualized as aggregate peak analysis (APA) plots using the coolpup.py suite (v0.9.5) (Flyamer et al. 2020). APA plots display the average contact signal centered on loop focal points, where the central pixel corresponds to the bin containing the differential or TF-specific chromatin contact. The resolution of the central bin and the total size of the displayed region, including the flanking sequences, vary by plot and are specified in the corresponding figure legends. The score displayed in the upper-left corner of each APA plot represents the enrichment of the central pixel intensity relative to the background.

#### h Drosophila chromatin state integration (Micro-C)

Micro-C differential chromatin interaction sets were integrated with the *Drosophila* S2 9-state chromatin model (Kharchenko et al. 2010) to annotate loop anchors using the *pybedtools* suite (v0.11.0) (Dale et al. 2011; Quinlan and Hall 2010). Two distinct approaches were used for this integration.

In the first approach (Figure 1), differential chromatin interactions in BEDPE format were overlapped with chromatin states in BED format using the *pairToBed* tool with -type either (default parameters), such that a chromatin state was considered associated with a given interaction if either anchor overlapped the state. Under this scheme, a single interaction could appear multiple times if its anchors overlapped with more than one chromatin state. This approach was used to determine the overall proportion of differential interactions overlapping each genome-wide chromatin state, as well as to characterize loop size distributions for interaction subsets associated with specific chromatin states.

In the second approach (Figure 4), both anchors of each chromatin interaction were independently annotated with their overlapping chromatin states using a custom Python script, generating chromatin state pairs for each interaction. This enabled analysis of the frequency and loop characteristics specific to each pairwise combination of chromatin states. As in the first approach, an interaction could appear multiple times if its anchors mapped to more than one chromatin state pair.

#### i Known and de novo motif discovery

Known and de novo motif discovery was performed at loci of interest using the HOMER motif discovery toolkit (v5.1) (Heinz et al. 2010). Specifically, the *findMotifsGenome.pl* script was run against the *Drosophila* dm6 genome assembly with the following parameters: -size 1000 -len 8,10,12,14,16,20 -mis 4 -S 10, specifying a 1kb window centered on each locus, motif lengths of 8–20bp, up to 4 mismatches, and a maximum of 10 motifs returned. The specific input loci varied by analysis and are detailed in the corresponding figure legends.

#### j roX2-RDD chromatin loop integration

Prior to integrating our Micro-C differential chromatin interaction sets with the *roX2*-RDD chromatin loops (<u>GSE207720</u>) produced by Tian et al., we normalized anchor bin sizes across all datasets to a uniform 5kb resolution using custom Python scripts. This normalization was necessary because the RDD method generates chromatin loops with variably sized anchors, which would otherwise confound direct comparison with our fixed-resolution Micro-C-derived loops. Although normalization reduced the total number of roX2-associated chromatin loops, the remaining set was sufficiently robust for downstream analysis. Complete loop counts for all datasets after normalization, including differential CLAMP- and GAF-dependent loops and *roX2*-RDD loops, are reported in Table S2.

#### k Comprehensive multi-omics visualization

Contact matrices, chromatin loops, bigWig signal tracks, and loci of interest were visualized using the CoolBox API (v0.3.9) (Xu et al. 2021). The following publicly available datasets were incorporated for visualization: H3K36me3 and H4K16ac ChIP-seq data from <u>GSE29537</u>, and MSL2 ChIP-seq data from <u>GSE37864</u>.

## Supporting information

Supplemental Statistics

## Data access

Micro-C and Hi-ChIP data is freely accessible via the Gene Expression Omnibus (GSE322508, GSE322509).

## Acknowledgements

We are grateful for the HHMI Gilliam award to J.A., the Blavatnik Family Fellowship to K.C., and the NIH R35 GM153389 award to E.L.

## Author contributions

J.A. and K.C. performed experiments, computational analysis, and wrote the manuscript. M.A. and L.C.S.A. and contributed to computational analysis. M.A., A.A, C.G., and M.R. contributed to cell maintenance, Hi-ChIP experiments, and Micro-C experiments. M.W.S. and K.G. performed Western blot experiments and contributed to methods. E.L. advised on experimental design, analysis, and manuscript writing.

