## Supplemental Figures for "Differential chromatin looping regulated by two GA-binding transcription factors creates an X-specific chromatin environment for dosage compensation"

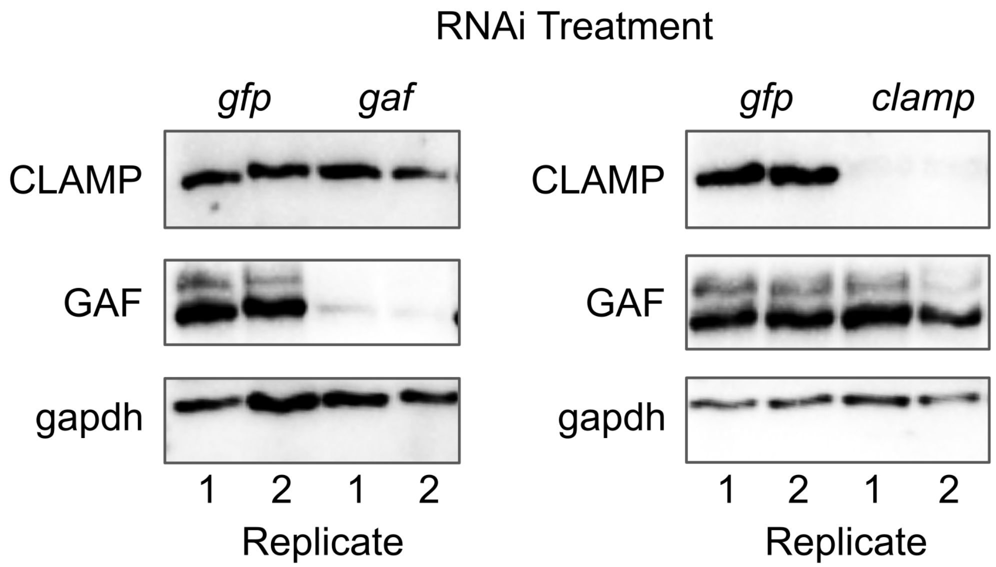


**Figure S1. Depletion of CLAMP or GAF in Micro-C samples.** Western blot analysis of protein extracts from cells treated with either *clamp* RNAi (*clampi*) or *gaf* RNAi (*gafi*). Blots were probed with both anti-CLAMP and anti-GAF antibodies across all conditions to confirm knockdown specificity. Treatment with *clampi* results in loss of CLAMP protein and treatment with *gafi* results in loss of GAF protein. anti-GAPDH antibody was used as a loading control to ensure equal amounts of protein lysate was loaded across all lanes.

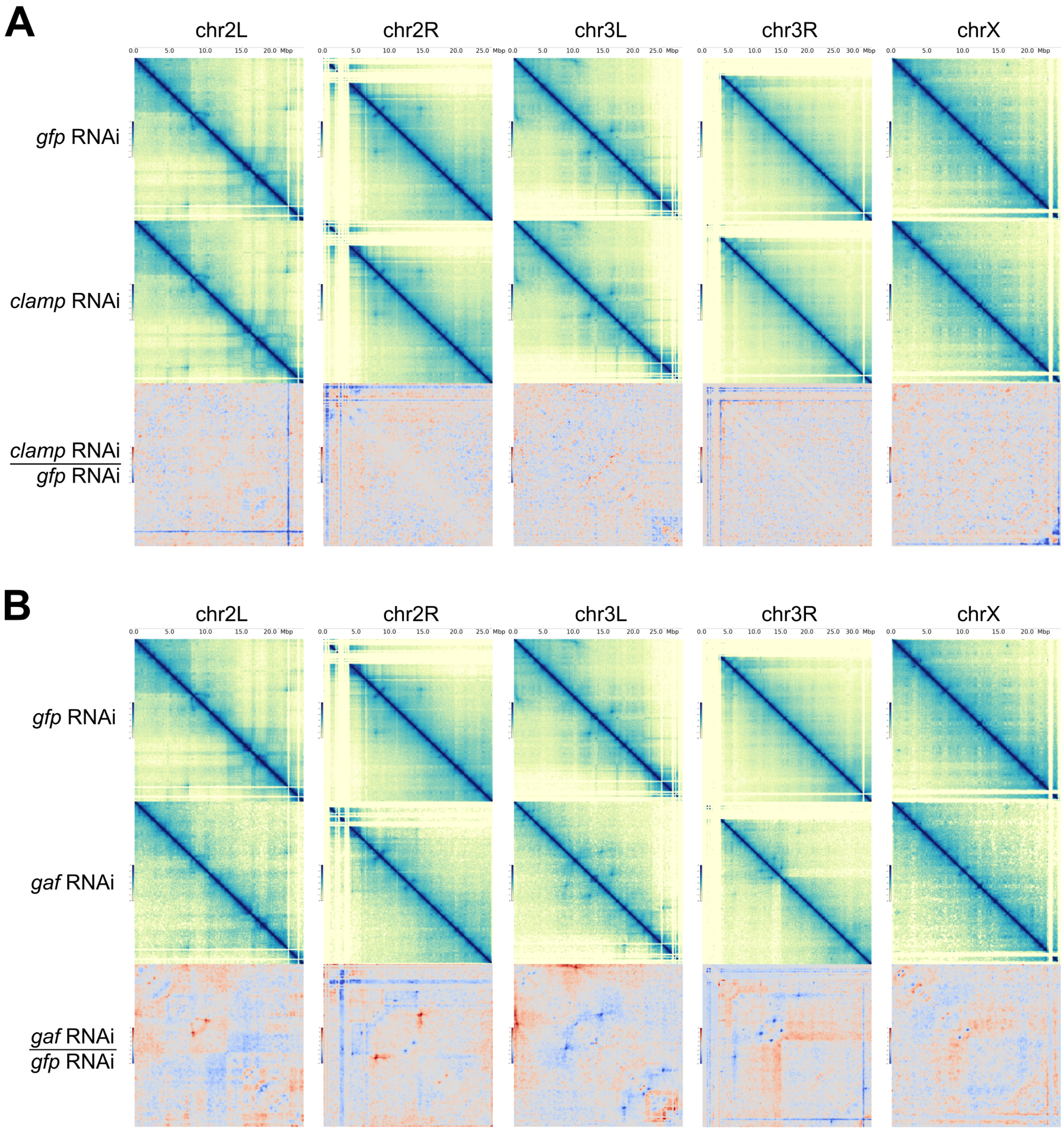


**Figure S2. Micro-C contact matrices across all *Drosophila melanogaster* chromosomes.** Micro-C control and post-RNAi treatment contact matrices are shown for all *D. melanogaster* chromosomes: chr2L, chr2R, chr3L, chr3R, and chrX. **(A) Differential Micro-C matrices for *clamp* RNAi. First row:** normalized contact maps for control *gfp* RNAi condition. **Second row:** normalized contact maps for *clamp* RNAi condition. **Third row:** differential contact matrices (*clamp* RNAi / *gfp* RNAi) where areas in red indicate increased contact frequency in *clamp* RNAi compared to *gfp* RNAi, while areas in blue indicate decreased contact frequency, with lighter colors representing smaller differences. (**B) Differential Micro-C matrices for *gaf* RNAi. First row:** normalized contact maps for the control *gfp* RNAi condition. **Second row:** normalized contact maps for *gaf* RNAi condition. **Third row:** differential contact matrices (*gaf* RNAi / *gfp* RNAi).

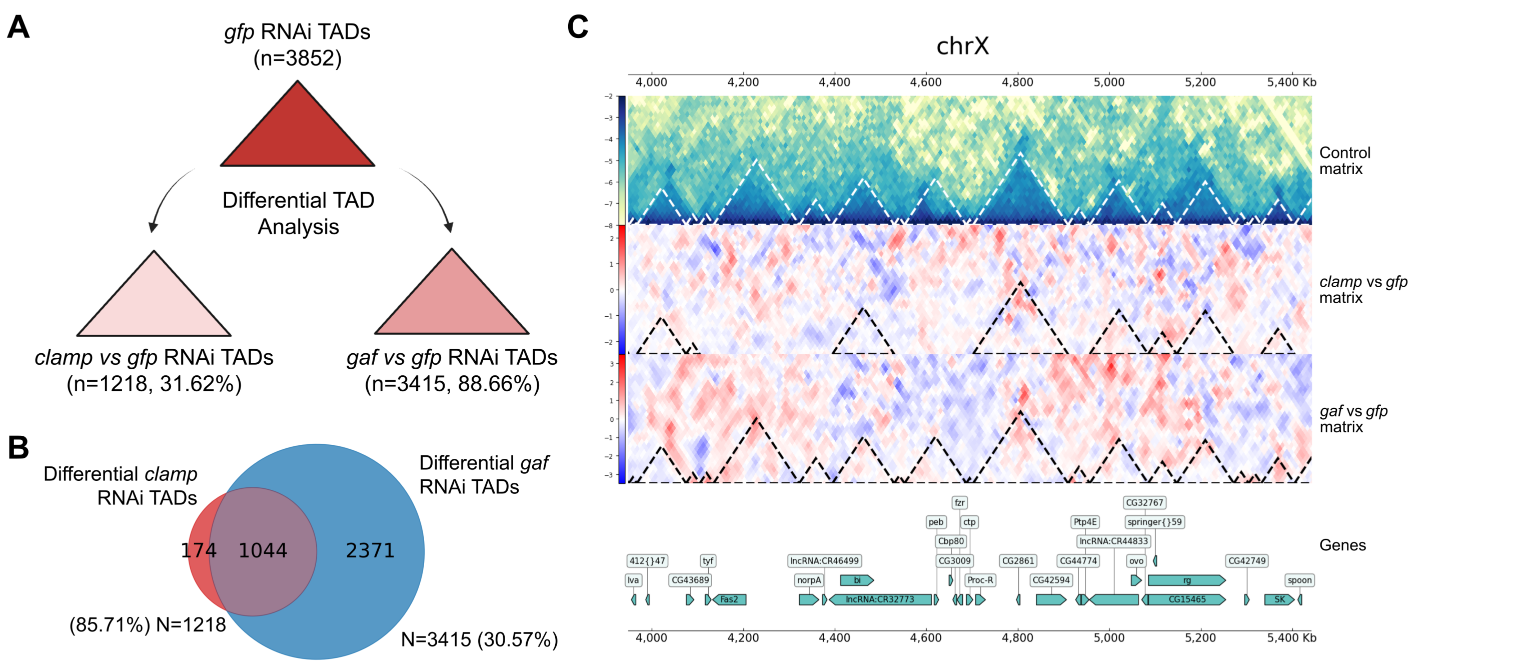


**Figure S3. TAD organization changes across the genome upon knockdown of CLAMP and GAF. (A) Quantification of *gfp* RNAi TADS and differential TAD comparison after treatment.** A total of n=3852 reference TADs were defined from the *gfp RNAi* control data (top triangle). A comparison to *clamp RNAi* (left) reveals n=1218 differential TADs, accounting for 31.62% of all reference TADs. A comparison to *gaf RNAi* (right) reveals n=3415 differential TADs, accounting for a larger 88.66% of reference TADs. This shows a greater genome-wide impact of *gaf* knockdown on TAD structure. **(B) Extensive Overlap of CLAMP- and GAF-Dependent TADs.** A Venn diagram illustrates that CLAMP and GAF share a large proportion of TAD changes following individual knockdown. Overlap between the sets of differential TADs identified for *clamp* RNAi (n=1218, red) and *gaf* RNAi (n=3415, blue) reveals a significant majority of CLAMP-dependent differential TADs (85.71%, N=1218) are also identified as GAF-dependent differential TADs. This overlap represents a subset (30.57%, N=3415) of the GAF-dependent differential TADs. **(C) Differential TADs on the X chromosome after *clamp* and *gaf* RNAi knockdown. Micro-C *gfp RNAi* control matrix:** White dashed lines indicate the positions of TAD boundaries with blue indicating greater contact and yellow indicating lower contact frequencies. ***Clamp* vs *gfp* matrix:** black dashed lines indicate differential TAD boundaries on the X chromosome after *clamp* RNAi treatment. ***Gaf* vs *gfp* matrix:** black dashed lines indicate differential TAD boundaries on X chromosome after *gaf* RNAi treatment, showing *gaf* knockdown leads to significantly different TADs across the 1,400kb window on X chromosome. **Genes:** Track of dm6 *Drosophila* X-chromosome genes across the same 1,400kb window.

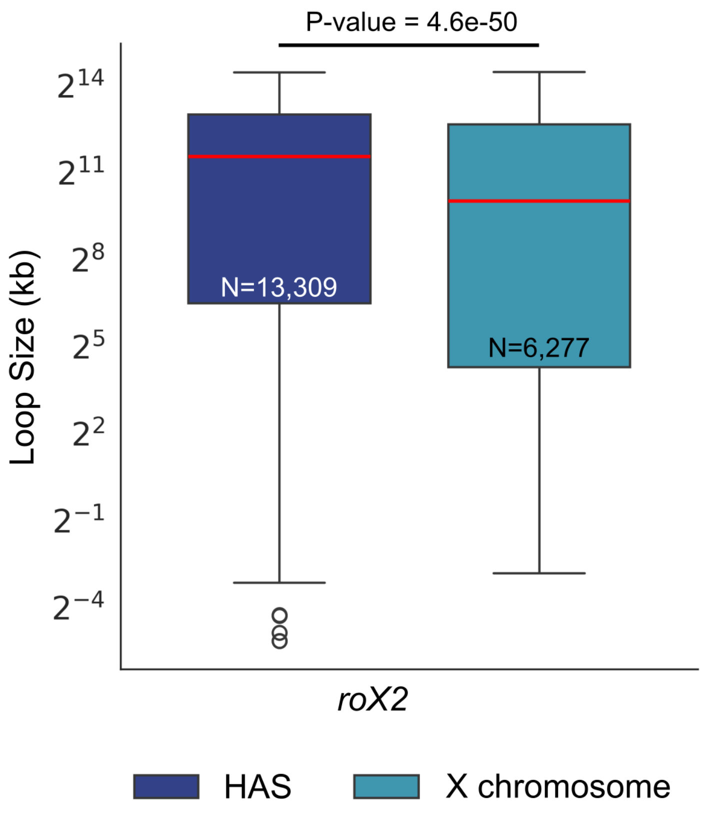


**Figure S4. The sizes of *roX2*-associated chromatin loops are significantly larger at HAS than on the X chromosome.** Box plot comparing the distribution of loop sizes (kb) at High Affinity Sites (HAS, dark blue, N=13,309) versus the X chromosome (light blue, N=6,277). Median size (red lines) of *roX2*-associated chromatin loops is significantly larger at HAS than non-HAS loops on X chromosome ($P=4.6 \times 10^{-50}$), as determined by a Mann-Whitney Test.

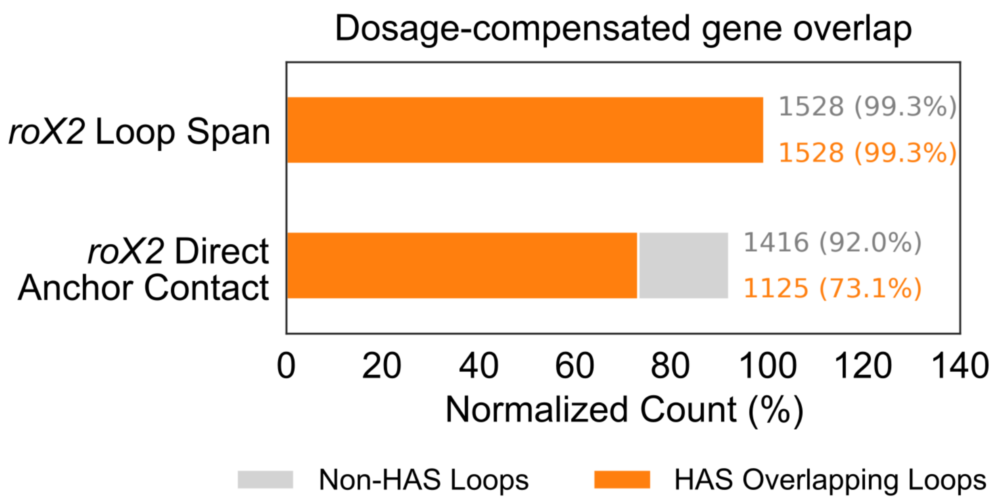


**Figure S5. Majority of *roX2*-associated chromatin loops encompassing dosage compensated genes are at HAS on the X chromosome.** Stacked bar plot quantifying the overlap between dosage-compensated genes on the X chromosome and two categories of *roX2*-associated Micro-C loops: **HAS Overlapping Loops** (orange) and **Non-HAS Loops** (gray). **Assessment of DC genes located within chromatin loop spans** associated with *roX2* versus DC genes directly at *roX2*-associated chromatin loop anchors. Nearly all identified dosage-compensated genes in *roX2*-associated chromatin loops (**99.3%**, n=1528) overlap with HAS. A high proportion (**73.1%**, n=1125) of dosage-compensated genes are located directly at HAS-overlapping loop anchors, compared to only **18.9%** (the remaining portion of the 92.0% total) found at anchors of loops not associated with an HAS.
